# Deep Learning-Based Unlearning of Dataset Bias for MRI Harmonisation and Confound Removal

**DOI:** 10.1101/2020.10.09.332973

**Authors:** Nicola K. Dinsdale, Mark Jenkinson, Ana I. L. Namburete

**Author notes:** Indicates Equal Contribution (Mark Jenkinson, Ana I. L. Namburete).

## Abstract

Increasingly large MRI neuroimaging datasets are becoming available, including many highly multi-site multi-scanner datasets. Combining the data from the different scanners is vital for increased statistical power; however, this leads to an increase in variance due to nonbiological factors such as the differences in acquisition protocols and hardware, which can mask signals of interest.

We propose a deep learning based training scheme, inspired by domain adaptation techniques, which uses an iterative update approach to aim to create scanner-invariant features while simultaneously maintaining performance on the main task of interest, thus reducing the influence of scanner on network predictions. We demonstrate the framework for regression, classification and segmentation tasks with two different network architectures.

We show that not only can the framework harmonise many-site datasets but it can also adapt to many data scenarios, including biased datasets and limited training labels. Finally, we show that the framework can be extended for the removal of other known confounds in addition to scanner. The overall framework is therefore flexible and should be applicable to a wide range of neuroimaging studies.

**Highlights:**

- We demonstrate a flexible deep-learning-based harmonisation framework
- Applied to age prediction and segmentation tasks in a range of datasets
- Scanner information is removed, maintaining performance and improving generalisability
- The framework can be used with any feedforward network architecture
- It successfully removes additional confounds and works with varied distributions

## 2. Introduction

The ability to combine datasets in neuroimaging between scanners and protocols is vital to achieve higher statistical power and is especially important when studying neurological conditions where limited data is available. Although some large scale neuroimaging projects now exist, such as the UK Biobank [43], most studies remain small and many of the larger studies are multi-site, such as ABIDE [11] and ADNI [25]. Pooling data across scanners and sites leads to an undesirable increase in non-biological variance, even when attempts have been made to harmonise acquisition protocols and use identical phantoms across imaging sites [53]. Multiple studies have confirmed this variation caused by scanner and acquisition differences including scanner manufacturer [22, 46], scanner upgrade [22], scanner drift [45], scanner strength [22], and gradient nonlinearities [27]. The ability to identify the presence of this variability has been further confirmed by several works applying the *‘Name the Dataset’* game [48] to neuroimaging datasets [20, 51].

The removal of this scanner-induced variance is therefore vital for many neuroimaging studies. Many existing methods are based on ComBat [26], an empirical Bayes method originally developed to remove ‘batch effects’ in genetics, which has been applied for harmonising values derived from structural [13, 38, 51], diffusion [14] and functional MRI [53], succesfully removing non-biological variance while preserving biological associations. ComBat uses multivariate linear mixed effects regression to account for both the biological variables and the scanner, to allow the modelling of the imaging features. For each site, a site-specific scaling factor *δ* is calculated, yielding a model that adjusts for additive and multiplicative effects. Furthermore, ComBat uses empirical Bayes to learn the model parameters, which assumes that model parameters across features are drawn from the same distribution; this improves the estimation of the parameters where only small sample sizes are available. ComBat has been further developed to include a term explicitly to model for variables of interest to preserve after harmonisation [51], model covariances [8], the incorporation of a generalized additive model (GAM) into the model, extending it to include nonlinear variations [38], and longitudinal studies [3]. ComBat, however, is usually applied to the harmonisation of image-derived values and defined associations. Random forests have also been used to approach the problem similarly, with the approach demonstrated for harmonising structural ROIs [19]. Other models also approach the harmonisation task through finding scaling factors for the raw image values, for instance MICA [52], which estimates nonlinear transformations of the voxel intensity values.

Harmonisation has been explored more for diffusion MRI than other modalities [5, 6, 7, 14, 24, 35, 33, 34, 36, 42] with many methods using spherical harmonics to harmonise the data [33, 34, 35, 36, 42] which greatly limits the ability of these methods to be applied to other modalities. Moyer et al [36] adopt variational autoencoders to produce scanner-invariant representations of the data for diffusion MRI. These feature representations can then be used to reconstruct the input images so that they are minimally informative of their original collection site. This work shows that using deep learning techniques to create feature representations that are invariant to scanner presents a strong candidate for MRI data harmonisation.

In addition to the variational autoencoders used in [36], other generative models have been used to harmonise MRI, largely based on deep learning models including U-Net [39] based models and cycleGAN [10, 56] based models [55]. These are limited by needing either paired or ‘travelling heads’ data for training for each site, which is expensive and infeasible to acquire in large numbers, but hard to evaluate without them [36]. With generative methods it is also very difficult to validate the generated ‘harmonised’ images and so there is a risk of unknown errors propagating through pipelines and affecting the results of any completed analysis; furthermore, there is very little exploration of this in the literature. In addition, these methods are data hungry and difficult to train, which leads to many of these methods being implemented in 2D [10] or on a patchwise basis [36]. Such implementations prevent the CNN learning context from adjacent slices or patches and can lead to errors in the reconstructed images, so that optimal performance is usually only achieved when the whole image can be scanned in a single forward pass [29].

The key measure of success for harmonisation methods is to be discriminative for the biological variable of interest whilst being indistinguishable with respect to the scanner used to acquire the data. Following the framework introduced by Ben-David et al [4], domain adaptation routinely concerns the scenario where we have a source domain with a large set of labels and a target domain with either no labels or a low number of labels. For this work, we consider only the case where the task to be performed is identical across domains. We would expect that these two domains would be related but not identical: for instance, the source and target domains could indicate different scanners or different acquisition protocols. The difference between the two domains means that a neural network trained on the source data would have a reduced performance on the target domain [50]; this degree of difference is known as *domain shift*[47]. The smaller the degree of the domain shift, the more likely the domain adaptation is to be successful. Domain adaptation techniques therefore attempt to find a feature space that performs a given main task whilst being invariant to the domain of the data. This is achieved by learning features using the source domain labels, but these features are transformed to a representation that should generalise to the target domain by simultaneously adapting the features to have the same distribution for both datasets. Therefore, domain adaptation should be applicable to the task of MRI data harmonisation, creating features that are indiscriminate with respect to scanner but discriminate with respect to the task of interest.

There have been many methods proposed for domain adaptation, the simplest of which use a divergence-based approach that tries to minimise some divergence criterion between feature distributions for the source and target data distributions, which should provide a domain-invariant representation. This relies on a feature representation existing, where the classifier is able to perform approximately equally on both tasks. Frequently used divergence measures include maximum mean discrepancy [40] that compares the means of two samples to determine if they are from the same distribution, and correlation alignment [44] that tries to align second-order statistics of the two distributions using a linear transformation. These methods are limited by the assumption that all scanner information can be removed by satisfying a simply definable condition, which is unlikely when dealing with a highly non-linear system such as MRI.

The similarity between domains is generalised further by using adversarial methods, where the features for each domain are forced to be identical for the family of possible nonlinear functions that can be produced by a given convolutional neural network. Adversarial domain adaptation was formalised with *Domain Adversarial Neural Networks* (DANNs) [18]. Based on the premise that for effective domain transfer to be achieved, predictions must be based on features that cannot discriminate between domains, they propose a generic framework that jointly optimises the underlying features as well as two discriminative classifiers that operate on these features. The first is a label predictor that predicts the class labels and the second is a domain classifier that aims to predict the source of the data. The overall aim of the network is then to minimise the loss of the label predictor and to maximise the loss of the domain classifier, such that the learnt features are discriminative for the main task but are completely unable to discriminate between domains. They show that this behaviour can be achieved for feedforward neural networks simply through the addition of a domain classifier formed of standard convolutional layers and a gradient reversal layer [17]. The gradient reversal layer allows the maximisation of the loss of the domain classifier whilst minimising the loss for the main task. Placed between the feature extractor and domain classifier, the gradient reversal layer acts as an identity function in the forward step, and during the backward pass, it multiplies the gradient function by −λ, where λ is a hyperparameter set empirically for a given experimental setup [18]. The extension of domain adaptation to *N* source domains – key to enable the harmonisation of more than two scanners – was formalised in [4] and demonstrated for adversarial domain adaptation in [55].

Another method for adversarial domain adaptation was introduced in [49] where, rather than using a gradient reversal layer to update the domain classifier in opposition to the task, they used an iterative training scheme. By alternating between learning the best domain classifier for a given feature representation, and then minimising a confusion loss which aims to force the domain predictions to be closer to a uniform distribution, they obtain a domain classifier that is maximally confused [49]. Compared to DANN-style unlearning networks, it is better at ensuring a domain classifier that is equally uninformative across the domains [2] because the confusion loss tries to force all softmax output values to be equal, which is vital for successful data harmonisation, especially as we extend to a larger number of source scanners. This work was then applied in [2] to facial recognition, not for domain adaptation, but for the explicit removal of identified sources of bias in their data, showing that the framework can be extended to ‘unlearn’ multiple factors simultaneously.

In this work we will show that the adversarial framework proposed in [49] can be adapted for use in harmonisation for deep learning tasks. By considering harmonisation to be a multiple source, joint domain adaptation task, we will show that we can produce shared feature representations that are invariant to the acquisition scanner while still completing the main task of interest across scanners and acquisition protocols with negligible performance compromise. Further, we will consider a range of likely data scenarios, such as the effect of having limited amounts of training data and the effect of having different distributions of data for different scanners, and show that the training framework is simply adapted to deal with these additional challenges. Finally, we will show that the framework can be used to remove other confounds in addition to harmonising for scanner. Whilst we demonstrate the network on age prediction and tissue segmentation, the framework is applicable to all feedforward networks and tasks and so should be applicable to the harmonisation of data across a wide variety of neuroimaging studies.

## 3. Methods

Before we explain the method, we will introduce some notation which will be used in the equations that follow. The network takes as input a set of MRI scans, ***X***, and outputs the predicted labels 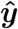, which can take the form of a vector of scalar values (for age regression) or class probability maps (for segmentation). We also have target labels ***y*** for the task of interest (regression or segmentation), which will be made available to the network during training. These data come from *N* distinct scanners or acquisition protocols, and for each scanner we have *S_n_* subjects, where *n*∈ {1..*N*}. Note that images acquired on the same *physical* scanner but using different acquisition parameters (eg. a change in T1 acquisition protocol) are treated as if they were acquired on different scanners.

We train a network formed of three parts, illustrated in Fig.1: a *feature extractor* with parameters **Θ**_*repr*_, a *label predictor* with parameters **Θ**_*p*_, and the *domain classifier* with parameters **Θ**_*d*_. The first two jointly form the network needed to perform the main task: the feature predictor takes the input image ***X*** and outputs a fully connected layer. This fully connected layer is then passed to the label predictor and outputs the main task label, ***y***. The domain classifier is also added to the output of the feature extractor to allow us to adversarially remove the scanner information. It therefore takes in the fully connected layer from the feature extractor and outputs softmax values where *p_n_* is the softmax value for the *n^th^* scanner and ***d*** is the true domain of the data corresponding to the acquisition scanner. We assume that we are able to obtain the domain label (e.g. scanner or acquisition protocol) for all available data points, which will be true for nearly all cases. We may, however, not have main task labels **y** (e.g. image annotations or subject age) available for all cases.

**Figure 1:**
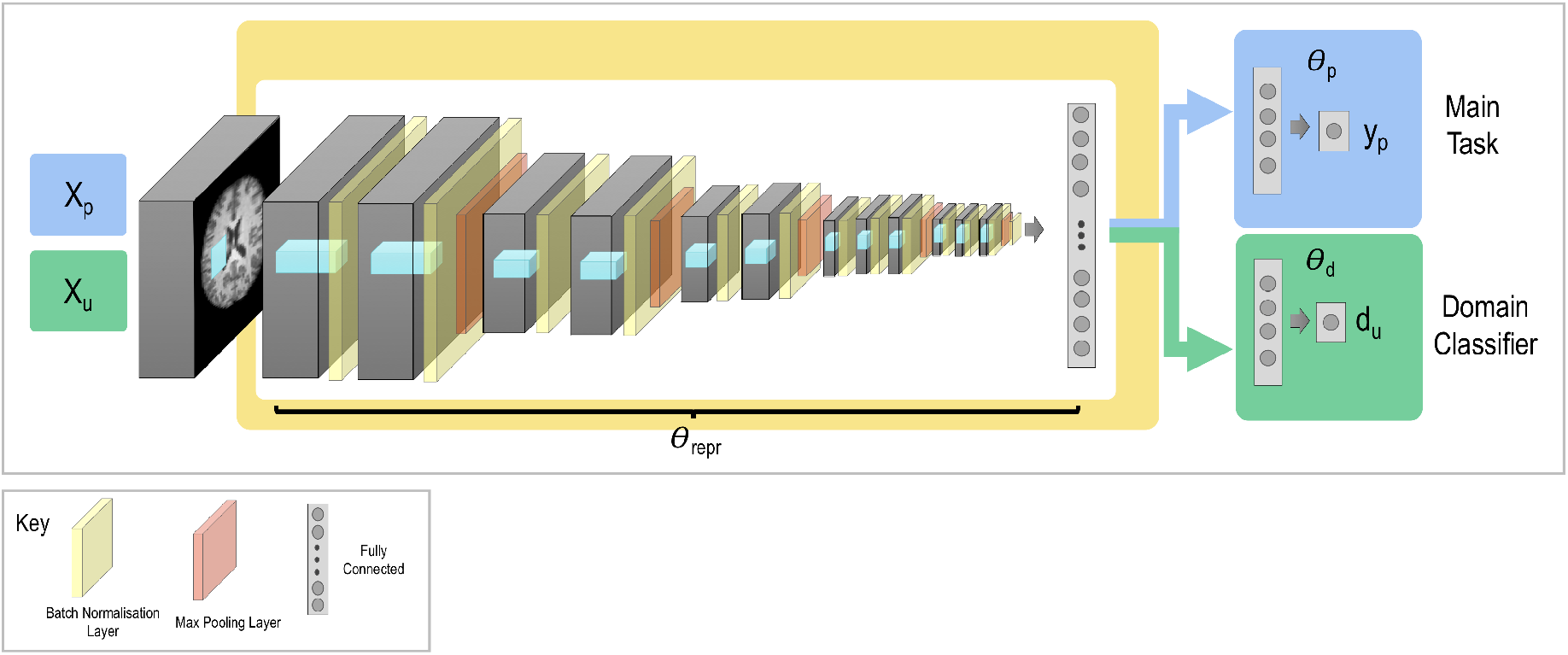
General network architecture. The network is formed of three sections: the feature extractor with parameters **Θ**_*repr*_, the label predictor with parameters **Θ**_*p*_, and the domain classifier with parameters **Θ**_*d*_. **X**_*p*_ represents the input data used to train the main task with labels **y**_*p*_, and **X**_*u*_ represents the input data used to train the steps involved in unlearning scanner with labels **d**.

Training of the network is controlled by optimising three loss functions following three consecutive steps in each training batch. Specifically, each loss function is optimised in turn, which results in three training iterations per batch. These work together to create a feature representation ***Q**_repr_* = *f*(**X**, **Θ**_*repr*_) – the activations at the final layer of the feature extractor – which we aim to make invariant to the scanner used for acquisition but discriminative for the main task of interest (e.g. segmentation or regression).

The first loss function pertains to the primary task of interest. For instance, for segmentation it may take the form of the Dice loss, whereas for a classification task it may be categorical cross entropy. This loss function is evaluated separately for each scanner (or dataset acquired with a different protocol) such that the optimisation is not driven by the largest dataset:

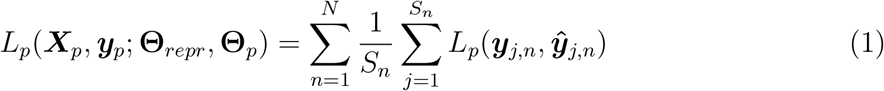

where ***X**_p_*⊂***X*** and *p* selects the subset of data for which main task labels are available, ***y**_p_*. *L_p_* is then the primary loss function being evaluated for the subjects acquired on scanner *n*∈ {1..*N*} and 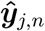 is the predicted label for the main task for the *j^th^* subject from the *n^th^* scanner. The notation *L_p_*(***X**_p_*, ***y**_p_*; **Θ**_*repr*_, **Θ**_*p*_) means that we evaluate the loss function for the data {***X**_p_*, ***y**_p_*}, which are fixed, given the current values of the parameters **Θ**_*repr*_, and **Θ**_*p*_ found by the optimisation process. It can therefore be seen that the optimizer controlling the training relating to this loss function is given access to the parameters in the feature extractor and the label predictor, as these are the values which are indicated to vary. The domain classifier is not involved in this stage.

The scanner information is then ‘unlearned’ (i.e. removed from the internal representation) using two loss functions in combination. The first is the domain loss that assesses how much scanner information remains in ***Q**_repr_*. It simply takes the form of categorical cross-entropy:

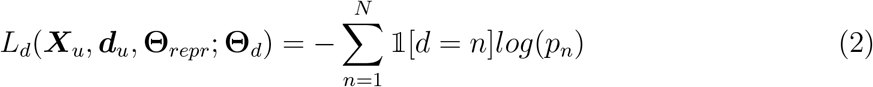

where *p_n_* are the softmax outputs of the domain classifier, and ***X**_u_*⊂***X*** and ***d**_u_*⊂***d*** where *u* indicates the subset of data to be used for unlearning. Although we assume that we are able to access domain labels for all of the data points, we do not necessarily use the full set for unlearning, depending on the scenario. Note the negative sign. This will be explored in the later experiments.

For this stage, the optimiser has access to only the weights in the domain classifier and has no influence on **Θ**_*repr*_, which is fixed, as indicated by the loss function *L_d_*(***X**_u_, **d**_u_*, **Θ**_*repr*_; **Θ**_*d*_). We therefore find the best domain classifier given the fixed feature representation and, hence, an understanding of the amount of scanner information remaining.

The second loss function controlling the unlearning is the confusion loss. Minimising this loss function aims to tune the parameters in the feature extractor **Θ**_*repr*_ such that all scanner information is removed from **Q**_*repr*_, thus, making it scanner invariant. When this feature representation is entirely scanner invariant, even the best domain classifier will be unable to predict which scanner the data used for acquisition, and so the softmax outputs of the domain classifier will all be equal, corresponding to random chance. The confusion loss therefore has the form:

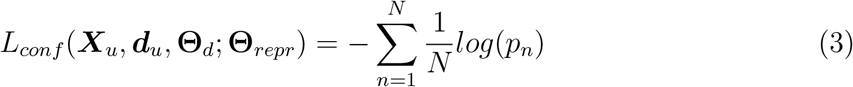

where ***X**_u_* and ***d**_u_* are the same subsets used in the previous loss function. This step only updates the features in **Θ**_*repr*_ and depends on the fixed value of **Θ**_*d*_ as indicated by ***L**_conf_* (***X**_u_,**d**_u_*,**Θ**_*d*_; **Θ**_*repr*_).

Stages 2 and 3 should be considered to be a unit and therefore the order in which they are updated is fixed. The confusion loss is most effective at removing crucial information to confuse the domain classifier once it has become effective at finding and utilising any domain information. Hence it is best to update the domain classifier first, to enable it to learn the domain information, prior to using the confusion loss. To this end, we must update equation (2) before equation (3).

Therefore, the overall method minimises the total loss function:

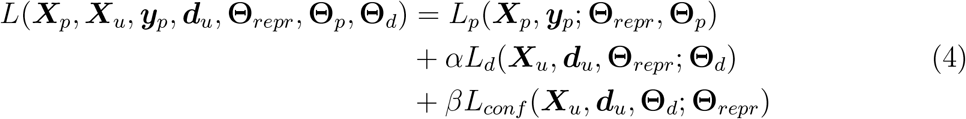

where *α* and *β* represent the weights of the relative contributions of the different loss functions.

Equations (2) and (3) cannot be optimised in a single step because they act in direct opposition to each other, hence the iterative update scheme and three forward/backwards passes per stage for each batch. This is not too computationally expensive as in each stage we only update a subset of the total parameters.

The domain classifier can only be used to assess how well domain information is being removed if it is capable of accurately predicting the domain *prior* to unlearning. In order to ensure this, we pretrain the network using equations (1) and (2) until the primary task reaches convergence. Provided that the domain classifier is able to accurately predict the scanner label at this stage, optimising the confusion loss in equation (3) such that the domain classifier performs no better than random chance corresponds to removing the scanner information in ***Q**_repr_*. In addition, this will decrease the number of epochs required for unlearning. As these are more computationally expensive, this serves to decrease the overall training time.

As shown by the loss functions, different sets of data can be used to evaluate the different loss functions. This not only means that we can unlearn scanner information for data points for which we do not have main task labels but also that we can unlearn scanner information using curated (or ‘matched’) subsets of data points when potentially problematic biases exist in the data as a whole. These scenarios are explored in the experiments to follow.

### 3.1. Age Prediction Task

We first consider the task of brain age prediction [9, 15, 16] as an example task to demonstrate the framework: ***X*** takes the form of the T1-weighted MRI images and ***y*** the true biological age values.

For these experiments, an architecture similar to the VGG-16 [41] network was used. A batch size of 32 was used throughout, with each batch constrained to have at least one example from each dataset. This increases stability during training but requires oversampling of the smaller datasets. The main task loss (equation (1)) was the Mean Square Error (MSE) for each scanner. All experiments were completed on a V100 GPU and were implemented in Python (3.6) and Pytorch (1.0.1) [37].

#### 3.1.1. Datasets

We used three datasets for these experiments: UK Biobank [43] (Siemens Skyra 3T) preprocessed using the UK Biobank Pipeline [1] (5508 training, 1377 testing); healthy subjects from the OASIS dataset [32] (Siemens Telsa Vision 1.5T) at multiple longitudinal time points, split into training and testing sets at the subject levels (813 training, 217 testing), and healthy subjects from the Whitehall II study [12] (Siemens Magnetom Verio 3T) (452 training, 51 testing). The model was trained using five-fold cross validation with the training data split as 80% training 20% validation; the results are reported for the hold out test set.

The input images for all the three datasets were resampled, taking account of the original voxel size, to 128 × 128 × 128 voxels, each of size 1×1×1mm using spline interpolation and then every fourth slice was retained, leaving 32 slices in the z direction (axial slices). This then maintains the physical size of the object and makes datasets comparable in this way, at the price of some interpolation-related changes to the intensities. Note that such resampling was necessary in order to enable us to use all the datasets within the same model.

Every fourth slice was selected so as to maximise coverage across the whole brain whilst minimising the redundancy and allowing us to use a larger batch size and number of filters. The inputs were also normalised to have zero mean and unit standard deviation. The distributions of the data can be seen in Fig.2. The p-values from T-tests on the dataset pairs show that there is only a significant difference between the distribution of the UK Biobank and Whitehall data - UK Biobank and OASIS p=0.22, OASIS and Whitehall p=0.09, UK Biobank and Whitehall p=0.001.

**Figure 2:**
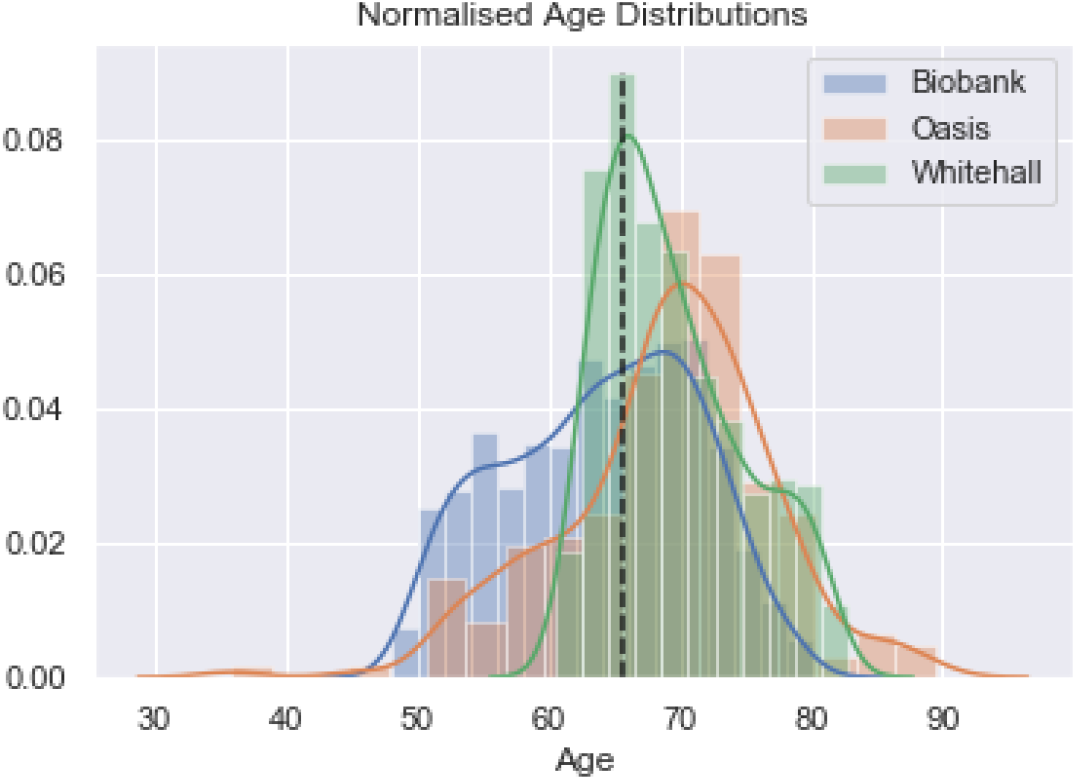
Normalised age distributions for the three datasets: Biobank, OASIS and Whitehall. The dashed line indicates the mean age of the three datasets.

#### 3.1.2. Basic Fully-Supervised Learning

We first consider the simplest training scenario where we have main task training labels available for all datasets being used and that all the datasets have similar distributions for the main task label. This means that there is not a high degree of correlation between the age and the scanner and we should, therefore, be able to remove scanner information from the feature representation without removing information that is discriminative for the age prediction task.

In this scenario, all three loss functions can be evaluated on a single combined dataset ***X*** where we have labels ***y*** for the full set and know the acquisition scanner for all data points ***d***, meaning that the overall method minimises the total loss function:

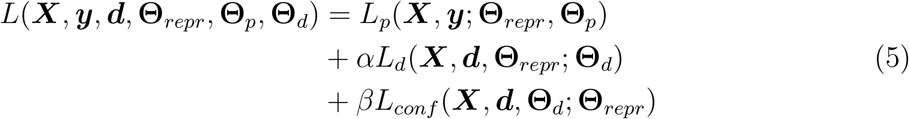

To allow comparison, we also train the network using normal training - training only the feature extractor and label predictor - on the different possible combinations of datasets, and compare to all combinations of datasets. As a form of ablation study, we also compare to standard training with the main task loss function evaluated as in equation (1). These comparisons are vital because we do not want the harmonisation process to significantly degrade the performance on the main task. We do not compare to existing harmonisation methods such as ComBat because these methods approach harmonisation differently to our method and so are not directly comparable.

#### 3.1.3. Biased Domains

When the distribution of the data for the main task is similar across all data points, unlearning can simply be completed on all of the data points. However, where there exists a large difference between the two domains, such that the main task label is highly indicative of the scanner, it is likely that the unlearning process will also remove information that is important for the main task. This latter scenario could, for instance, be where the age distributions for the two studies are only slightly overlapping or where nearly all subjects with a given condition are collected on one of the scanners.

To reduce this problem, we utilise the flexibility of the training framework, and, whilst evaluating the main task on the whole dataset, we perform the scanner unlearning (equations (2) and (3)) on only a subset of the data. For the case of age prediction, we perform unlearning only on the overlapping section of the data distributions. If we were to consider the case where we had data from both subjects and healthy controls and, for instance, most of the subjects had been scanned on one of the two scanners, we could perform unlearning on only the healthy controls rather than the whole dataset. As we do not need main task labels for the data used for unlearning, unlearning could be performed on a separate dataset so long as the scanner and protocol remained identical.

We subsample the Biobank and OASIS datasets so as to create three degrees of overlapping datasets: 5 years, 10 years and 15 years (Fig 3) and test on the same test sets as before, spanning the whole age range. We compare three training options: i) training normally on the full combination of the biased datasets, ii) unlearning on the whole distribution, and iii) unlearning only on the overlapping section of the distributions.

**Figure 3:**
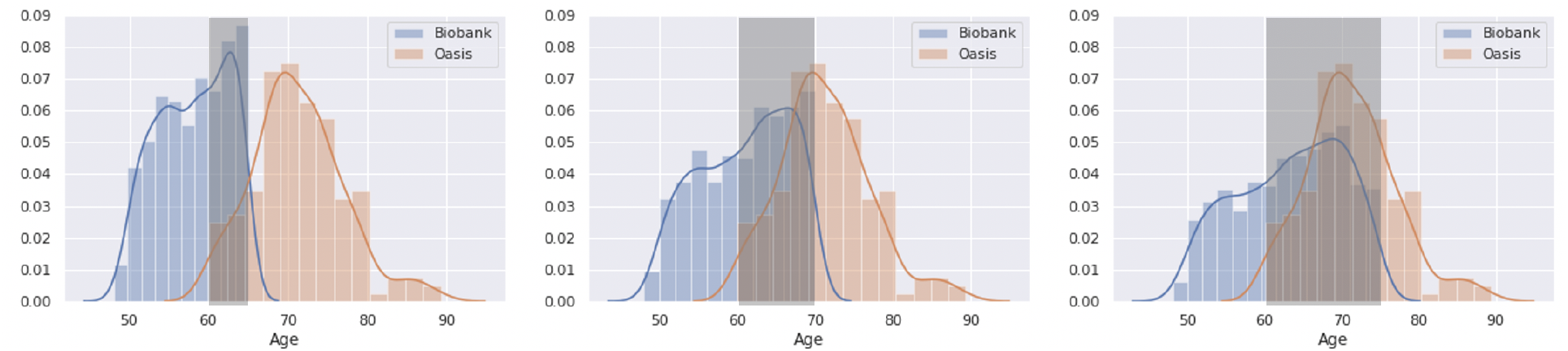
The three biased datasets used in this experiment with 5 years, 10 years and 15 years overlap. Only the shaded overlap region is used for unlearning scanner information; all the data points are used to evaluate the main loss function.

#### 3.1.4 Extension to ABIDE data

Having demonstrated the network on three scanners, we now demonstrate the network on a multi-site dataset, ABIDE [11]. We split the data into 90% training and 10% testing across each scanner and process as described in Section 3.1.2. We therefore have data from 16 scanners, including Philips, Siemens and GE scanners, each with between 27 and 165 scans per site, for training. This is representative of many neuroimaging studies, with relatively low numbers of scans available for each site. Following the same fully supervised framework, as demonstrated in Section 3.1.2, we compare training on the combination of data from all 16 sites, to the unlearning approach.

A batch size of 32 was used in training, with the batch constrained to have at least one example from each scanner. To achieve this, the smaller datasets were over sampled such that the number of batches was limited by the largest dataset. This was performed because it was found that having an example of each dataset in each batch led to higher stability during training. The over sampled data points were not augmented so that we could be sure improvement was not due to the augmentation, but, in practice, this would be a sensible step.

#### 3.1.5. Removal of other categorical confounds

In addition to unlearning scanner information to harmonise the data, we can also adapt the framework to explicitly remove other confounds. As shown in Fig 4, an additional data pair is added to training and an additional two loss functions are added to the overall loss function per confound. For each confound we wish to remove, we add a *confound classification loss*, which, like the domain classification loss, identifies how much information relating to the confound remains in the feature space, and a *confound confusion loss*, which aims to modify the feature space so as to remove the confound information by penalising deviations from a uniform distribution for the softmax predictions for the given task.

**Figure 4:**
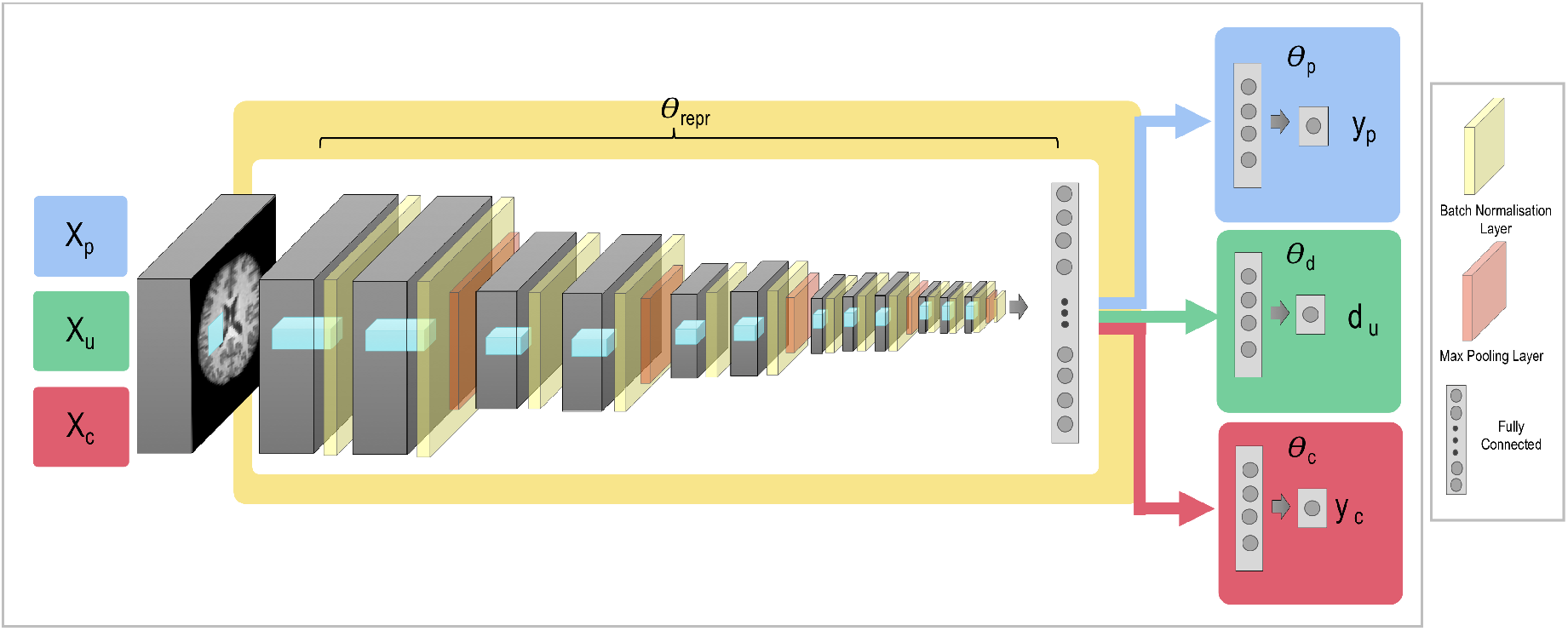
The general network architecture can be adapted to allow us to also remove other confounds from the data. In addition to the datasets used for the main task and the scanner unlearning, we add a data pair where ***X**_c_* are the input images used for deconfounding and ***y**_c_* are the confound labels.

Therefore, the overall method will minimise the loss:

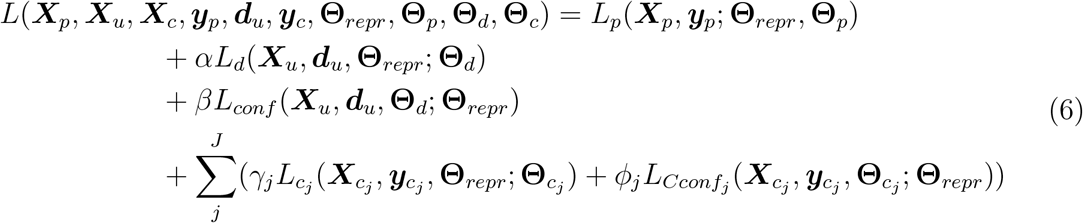

where we consider *J* different confounds we wish to remove, *γ_j_* is the weighting for the classification loss *L_c_j__* for the *j*^*th*^ confound, and similarly *ϕ_j_* is the weighting of the confound confusion loss *L_Cconf_j__* for the *j*^*th*^ confound. We demonstrate this with sex as the confound and so the classification loss can simply be the binary cross-entropy loss. If the confound to be unlearned took the form of a continuous variable, such as age, then following [2] it would need to be split into discrete bins.

We consider three different scenarios for removing sex as a confound while completing the age prediction task. We first consider the simplest case, where sex is approximately equally distributed across age and scanner. In this case we can simply evaluate all the loss functions for the whole set of input images – assuming all labels are available for input images for all tasks.

We also explore the scenarios of the confound being correlated with a) the scanner and b) the main task. For correlation with scanner, we create datasets where 80% of the subjects in the OASIS dataset are male and 80% of the subjects in the Biobank dataset are female. For correlation with age, we consider the case where 80% of the subjects under the median age across the two datasets (65) are female and 80% of the subjects over the median age are male. For both scenarios, we still test on the full test set, and we compare normal training, unlearning sex on the whole training set, and unlearning sex on a subset curated to be balanced with respect to the generated bias. In the last case, the loss functions for the main task and unlearning of scanner are evaluated across the whole datasets and only the equations controlling unlearning the sex confound are evaluated for the subsets.

#### 3.1.6. Removal of Continuous Confounds

While the approach for the removal of categorical confounds, such as sex, is clear from the section above, the extension to continuous variables is less clear. If we reframe our network such that sex prediction is now the main task, we can consider the process of removing the age information.

The continuous age labels could be approximately converted into categorical labels by binning the data into single-year bins spanning across the age range. This, however, would not encode the fact that a prediction of 65 for a true label of 66 encodes more true age information than a prediction of 20. We therefore convert the true age labels into a softmax label around the true age, normally distributed as a 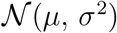 where *μ* was the true age label and *σ* was set to 10 empirically, allowing us to maintain relative information between bins. The value of 10 was chosen as, when used in normal training, this minimised the mean absolute error (MAE) on the test set. The loss function for the main task then becomes the Kullback-Leibler (KL) divergence between the true softmax distribution and the predicted softmax outputs. Unlearning can then be achieved as before, with the aim still being to make all the softmax outputs equal, so that no information about the age of the subject remains.

### 3.2. Segmentation Task

Having demonstrated the unlearning process on the age prediction task, we now consider the task of segmentation. We specifically consider the case where our network architecture takes the form of a U-Net [39], the most popular network for biomedical image segmentation, where the skip-connections, which form a crucial part of the architecture, have the possibility of increasing the complexity of unlearning. Again, the input images ***X*** are T1-weighted input images but the labels take the form of tissue segmentations (grey matter/white matter/CSF) produced using FSL FAST [54] as a proxy to manual segmentations, which were converted to one-hot labels for the segmentation. We consider examples from the UK Biobank [43] (2095 training, 937 testing) and healthy subjects from the OASIS dataset [32] (813 training, 217 testing). As before, these were resized to 128 × 128 × 128 voxels (each of which is 1 x 1 x 1 mm in size) using spline interpolation and the intensity values normalised, but then they were split into 2D slices so that we trained a 2D network. The labels were interpolated using trilinear interpolation and then thresholded at 0.5 to create categorical labels. Multi-class Dice Loss was used as the primary task loss function. All experiments were completed on a V100 GPU and were implemented in Python (3.6) and Pytorch (1.0.1).

#### 3.2.1 Basic Fully Supervised Learning

As can be seen in Fig.5, the general structure is identical to that used for age prediction, apart from the location of the domain classifier. In the case of the age prediction task, the unlearning was performed at the first fully connected layer, chosen as it was both a reduced representation of the data and because it was at the end of the network so there were few weights afterwards for the network to re-learn how to exploit any remaining scanner information were it not fully removed by the unlearning process. In the case of the U-Net architecture, the most compressed representation of the information is at the bottleneck (B) but there are both several upsampling layers and skip-connections after this point. Therefore, if we were to only complete unlearning at the bottleneck, all scanner information would most likely be re-learned by the subsequent layers in the upsampling branch and so might still influence the output segmentations.

**Figure 5:**
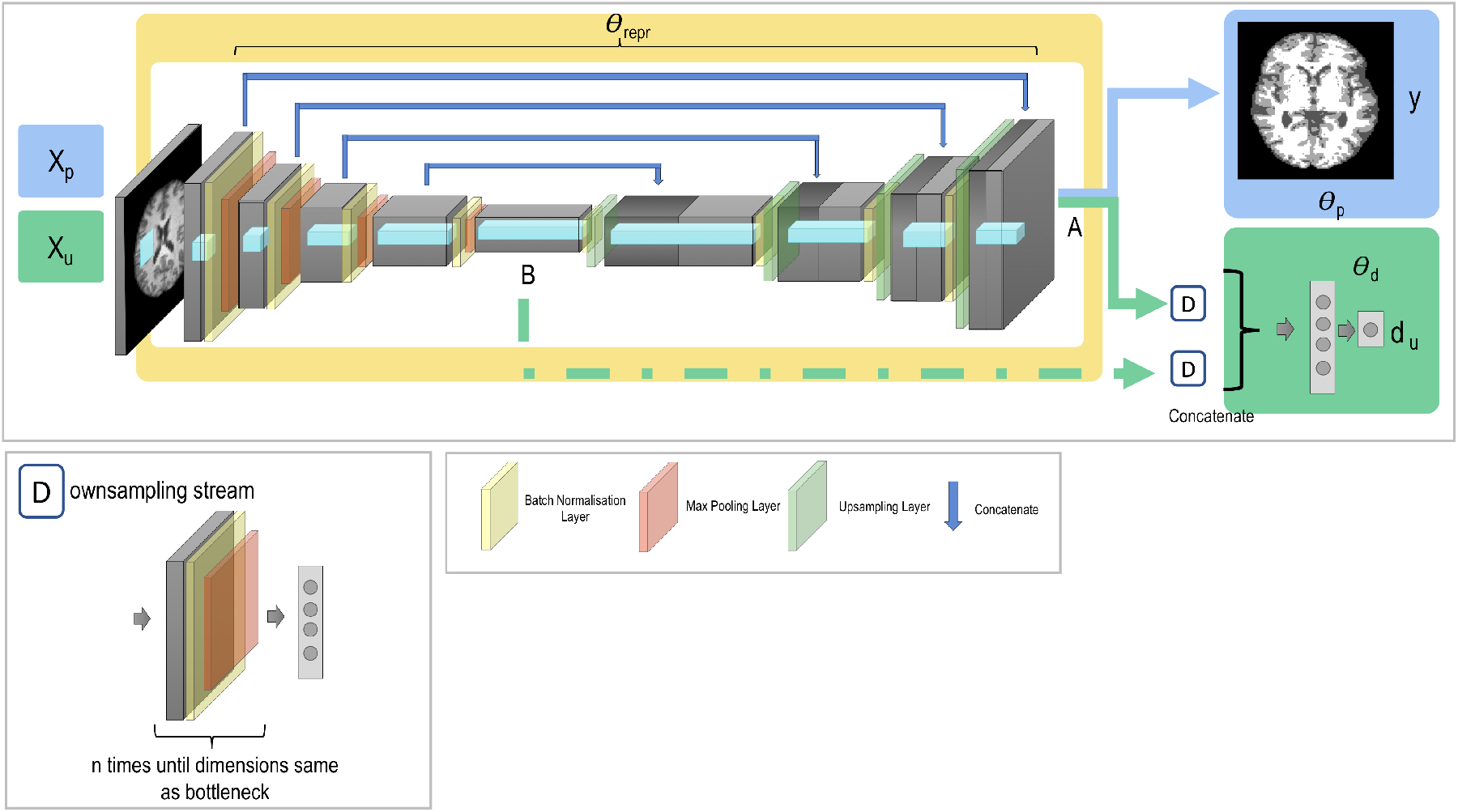
Network architecture used for unlearning with segmentation. ***X**_p_* represents the input images used to evaluate the primary task with **y** being the main task label segmentations. ***X**_u_* are the input images used for unlearning scanner information with domain labels *d_u_*. The domain discriminator for unlearning can be attached from A, B or the two in combination. If it is attached from A and B together, the first fully connected layers (the output of the two downsampling branches D) are concatenated together to produce a single feature representation.

We therefore consider unlearning from both the end of the network before the label predictor (A), the bottleneck (B) and the combination of the two, formed by concatenating the first fully-connected layers to form a single domain prediction. We also compare to standard training as a benchmark. We utilise the simplest training regime, where we have segmentation labels available for all the data from all scanners and that we have identical segmentation tasks. Therefore the overall loss function takes the same form as equation (5) and all of the loss functions are evaluated on all of the data points.

#### 3.2.2. Semi-Supervised Learning

Finally, we consider the scenario where we have very few or no labels available for one of the scanners – this is a very likely scenario for segmentation where manual labels are time consuming and difficult to obtain. We therefore assume access to one fully labelled dataset (UK Biobank) - and another dataset for which we do not have many labels. While the unlabelled data points cannot be used to evaluate the main task, they can be used for scanner unlearning.

No changes to the architecture are required; rather, we simply evaluate the main task for those data points for which we have main task labels and use all data points for unlearning such that the overall method minimises:

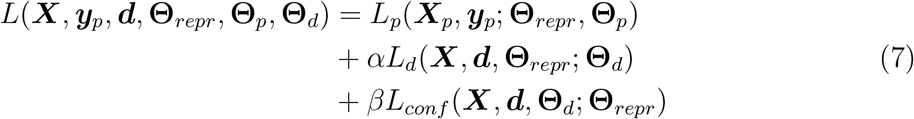

where ***X**_p_* is the subset of **X** for which we have main task labels ***y**_p_* available and the full dataset ***X*** is used in unlearning scanner information.

We explore the effect of increasing numbers of data points on the final segmentation, comparing to normal training on the combination of the full UK Biobank dataset and OASIS with available segmentations.

### 3.3. Ethics Statement

UK Biobank: has approval from the North West Multi-centre Research Ethics Committee (MREC) to obtain and disseminate data and samples from the participants, and these ethical regulations cover the work in this study. Written informed consent was obtained from all participants. Details can be found at www.ukbiobank.ac.uk/ethics.

Whitehall dataset: ethical approval was granted generically for the “Protocol for non-invasive magnetic resonance investigations in healthy volunteers” (MSD/IDREC/2010/P17.2) by the University of Oxford Central University / Medical Science Division Interdisciplinary Research Ethics Committee (CUREC/MSD-IDREC), who also approved the specific protocol: “Predicting MRI abnormalities with longitudinal data of the Whitehall II sub-study” (MSD-IDREC-C1-2011-71).

OASIS dataset: was previously collected under several study protocols at Washington University. All studies were approved by the University’s Institutional Review Board (IRB). All subjects gave written informed consent at the time of study participation. The University’s IRB also provided explicit approval for open sharing of the anonymized data. More information relating to this can be found in [32].

ABIDE dataset: prior to data contribution, sites are required to confirm that their local Institutional Review Board (IRB) or ethics committee have approved both the initial data collection and the retrospective sharing of a fully de-identified version of the datasets (i.e., after removal of the 18 protected health information identifiers including facial information from structural images as identified by the Health Insurance Portable and Accountability Act [HIPAA]). More details can be found in [11].

## 4. Results

### 4.1. Age Prediction Task

#### 4.1.1. Basic Fully-Supervised Learning

The results from training with all three datasets separately and the different combinations can be seen in Table 1. The scanner classification accuracy is achieved by taking the frozen feature representation ***Q**_repr_* from training, and training a classifier based on this. For the unlearning method, this is the domain classifier that was used during training but fine tuned to convergence. For normal training, an identical domain classifier is trained using the frozen feature representation as input until convergence. Therefore, the closer the value is to random chance, the less informative the feature representation is about scanner, thus meaning the influence of the scanner on the prediction is reduced. The results from training on all combinations of datasets both with standard training, and standard training using our loss function for the main task conditioned on each scanner, are shown for comparison.

**Table 1:**
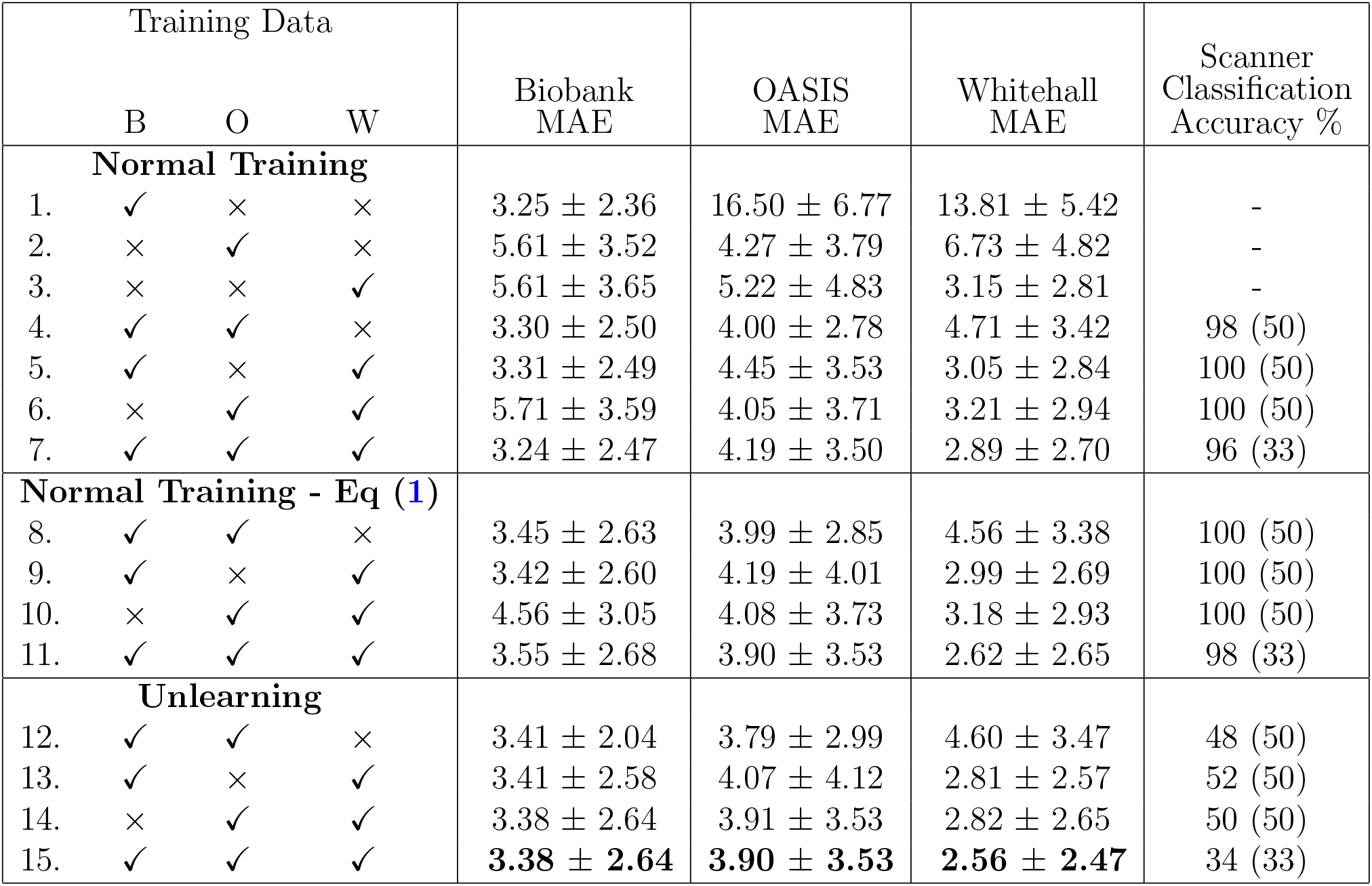
Results comparing unlearning to training the network in different combinations on the datasets. Mean absolute error is reported in years. Scanner accuracy is the accuracy achieved by a domain classifier given the fixed feature representation at convergence, evaluating only for the datasets the network was trained on. Number in brackets indicates random chance. B = Biobank, O = OASIS, W = Whitehall. p values can be found in the supplementary material. Bold indicates the experiment with the best average across the datasets.

In Fig.6, a T-SNE plot [31] can be seen, allowing visualisation of the activations at the output of the feature extractor, ***Q**_repr_*. It can be seen that, before unlearning, the features produced are entirely separable by scanner, but after unlearning the scanner features become jointly embedded and so the feature embedding is not informative of scanner.

**Figure 6:**
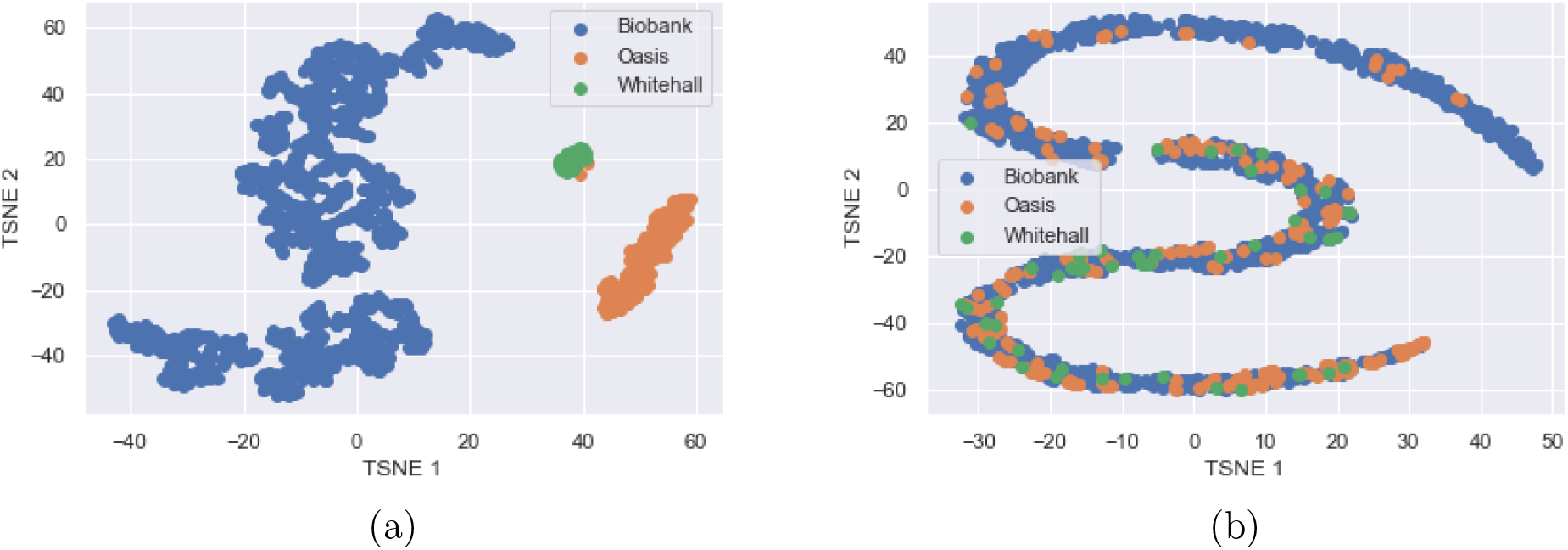
a) T-SNE plot of the activations of the fully connected layer, ***Q**_repr_*, before unlearning. It can be seen that the domains can be almost entirely separated, except for two data points grouped incorrectly, showing that data from each scanner has its own distinct distribution. b) T-SNE plot of the activations of the fully connected layer, ***Q**_repr_*, after unlearning. It can be seen that, through the unlearning, the distributions become entirely jointly embedded.

The T-SNE demonstrates that without unlearning, not only are there identifiable scanner effects but that they also affect the age predictions. On the other hand, it can be seen that we are able to remove scanner information using our unlearning technique, such that the data points for all three scanners share the same embedding. This is confirmed by the scanner classification accuracy being almost random chance after unlearning has been completed.

It can also be seen that unlearning does not decrease substantially the performance on the main task. Graphs showing the different loss functions with training can be found in the supplementary material. If we consider the case of training on all three datasets, we actually achieve an overall improvement in performance using unlearning across the three datasets (comparing lines 7 and 15 in Tab.1) with the performance on the OASIS and Whitehall datasets substantially improving. The performance on the UK Biobank dataset decreases, but this is as would be expected because the UK Biobank dataset has a much larger size and so, with normal training, the network is driven by this dataset whereas, with unlearning, all of the datasets drive the performance. Comparing to line 11, where the same loss function was used for the main task, evaluated separately for each dataset, but unlearning was not performed, we can see that there was an improved performance when the unlearning was added and so the unlearning itself provides actual improvement on the main task. It can also be seen that the scanner classification accuracy is almost perfect when using normal training (96%), or the loss function evaluated on the datasets separately (98%), and is random chance after unlearning (34%), clearly demonstrating that the unlearning process removes scanner information from the feature space.

It can also be seen that the unlearning process creates a feature space that generalises better to the third unseen dataset. If, for instance, we compare lines 5 and 13 we can see that not only is there an improved performance on the OASIS and Whitehall datasets used for training, but there is also a significant improvement in the performance on the UK Biobank dataset, which was unseen by the network. This, therefore, suggests that the unlearning process not only creates features that are invariant to the scanners that were present in the training set, but also that these features are more generalisable and so can be more effectively applied to other scanners. It can again be seen there is an improvement between normal training with our loss function evaluated on each scanner separately and the performance with unlearning (lines 9 and 13), showing that the increase in generalisability of the features is due to the unlearning process. The same pattern can be seen for training on the other possible pairs of datasets.

The generalisability of the features between datasets and the reduction of the scanner classification accuracy together demonstrate that the unlearning process successfully harmonises the differences between scanners and protocols.

The results were also found to be robust to the choice of hyperparameters. An exploration of this can be found in the supplementary materials.

We considered if it were better to use the full imbalanced datasets to maintain the number of data points or if this hindered the unlearning process. Therefore, we randomly downsampled the Biobank and OASIS datasets to have the same number of examples as the Whitehall dataset (452 subjects). Testing was evaluated on the same test dataset as in the previous experiments. The results can be seen in Table 2. For both normal training and unlearning, there was a large decrease in performance using the balanced datasets, but the decrease was less pronounced when using unlearning than when using normal training. Using balanced datasets made no difference to the ability to classify scanner: with normal training the classifier was still able fully to distinguish between scanners; with unlearning, the classification was almost random chance. Therefore, any advantage gained from having balanced datasets is outweighed by the reduction in performance from having reduced numbers of training examples and so we are better off training on the whole dataset. Given that the method is able to remove nearly all information with significant imbalance in this case (Biobank 5508 training data points compared to Whitehall’s 452) it is likely that unlearning would be sufficient in most cases.

**Table 2:**
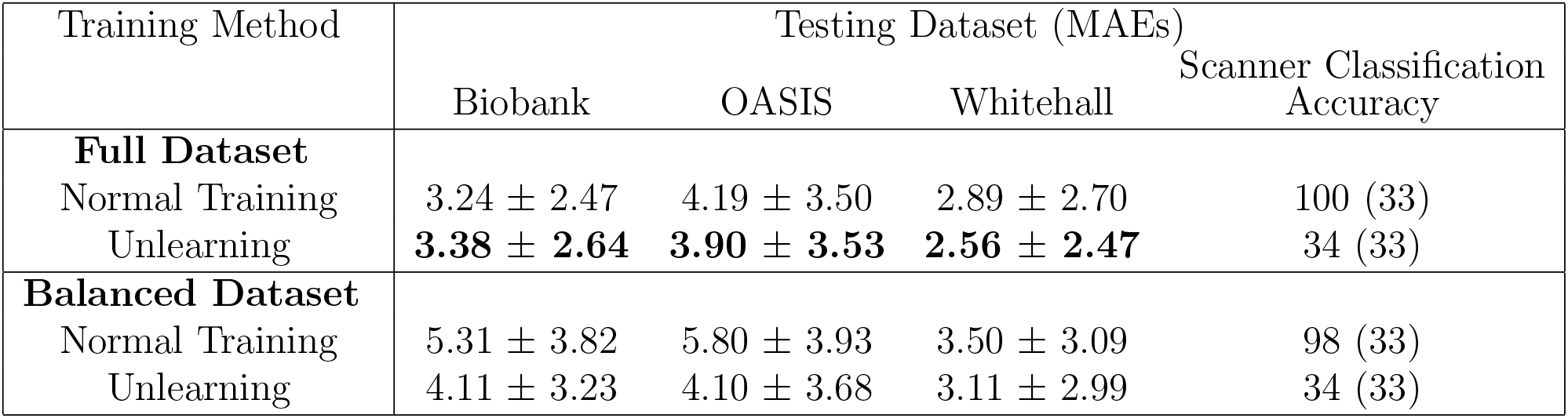
Comparison of the effect of training with the full datasets with imbalanced numbers from each scanner versus training with balanced numbers of subjects from each scanner (452 per scanner, randomly selected compared to 5508 Biobank, 813 OASIS and 452 Whitehall), comparing both normal training and unlearning. The method giving the best average across datasets is highlighted.

| Training Method | Testing Dataset (MAEs) |  |  | Scanner Classification Accuracy |
| --- | --- | --- | --- | --- |
|  | Biobank | OASIS | Whitehall |  |
| **Full Dataset** |  |  |  |  |
| Normal Training | $3.24 \pm 2.47$ | $4.19 \pm 3.50$ | $2.89 \pm 2.70$ | 100 (33) |
| Unlearning | **$3.38 \pm 2.64$** | **$3.90 \pm 3.53$** | **$2.56 \pm 2.47$** | 34 (33) |
| **Balanced Dataset** |  |  |  |  |
| Normal Training | $5.31 \pm 3.82$ | $5.80 \pm 3.93$ | $3.50 \pm 3.09$ | 98 (33) |
| Unlearning | $4.11 \pm 3.23$ | $4.10 \pm 3.68$ | $3.11 \pm 2.99$ | 34 (33) |

#### 4.1.2. Biased Datasets

We assessed the performance of the network training with biased datasets curated from subsets of the Biobank and OASIS datasets. We considered the cases of 5-year, 10-year and 15-year overlaps where the smaller the overlap, the harder the task. The trained networks are tested on the same test sets used above, with subjects across the whole age range included.

The results can be seen in Table 3 where we compare normal training, naïve unlearning on the whole dataset, and unlearning on only the overlapping age range. It can be seen that for all three degrees of overlap, standard training leads to much larger errors than unlearning and that unlearning on only the overlap gives a lower error than unlearning on all data points. Plots of the MAEs with age can be seen in Fig.7 for the 10-year overlap case. It can be seen that the overall lowest MAEs are achieved across the age range when unlearning is performed only on the overlapping subjects.

**Table 3:** MAE results for Biobank and OASIS data from training with datasets with varying degrees of overlap as shown in figure 3. Standard refers to training normally and naïve unlearning refers to unlearning on the whole of the datasets. Scanner accuracy is calculated by training a domain classifier on the fixed feature representation. Random chance is given in brackets. The method with the best average across the datasets for each degree of overlap is highlighted.

| Method | Biobank<br>MAE | OASIS<br>MAE | Scanner<br>Classification<br>Accuracy (%) |
| --- | --- | --- | --- |
| **5 Years Overlap** |  |  |  |
| 1. Standard | $16.5 \pm 5.94$ | $15.5 \pm 6.95$ | 100 (50) |
| 2. Naïve Unlearning | $6.11 \pm 3.99$ | $4.44 \pm 4.20$ | 58 (50) |
| 3. Unlearning on Overlap | **$5.49 \pm 3.67$** | **$4.37 \pm 4.05$** | 53 (50) |
| **10 Years Overlap** |  |  |  |
| 4. Standard | $9.66 \pm 5.83$ | $13.6 \pm 6.58$ | 100 (50) |
| 5. Naïve Unlearning | $4.20 \pm 2.90$ | $4.29 \pm 4.01$ | 56 (50) |
| 6. Unlearning on Overlap | **$3.93 \pm 2.81$** | **$4.04 \pm 3.86$** | 52 (50) |
| **15 Years Overlap** |  |  |  |
| 7. Standard | $8.91 \pm 5.31$ | $10.4 \pm 5.55$ | 100 (50) |
| 8. Naïve Unlearning | $3.82 \pm 2.84$ | $4.39 \pm 4.07$ | 57 (50) |
| 9. Unlearning on Overlap | **$3.75 \pm 2.78$** | **$3.99 \pm 3.52$** | 50 (50) |

**Figure 7:**
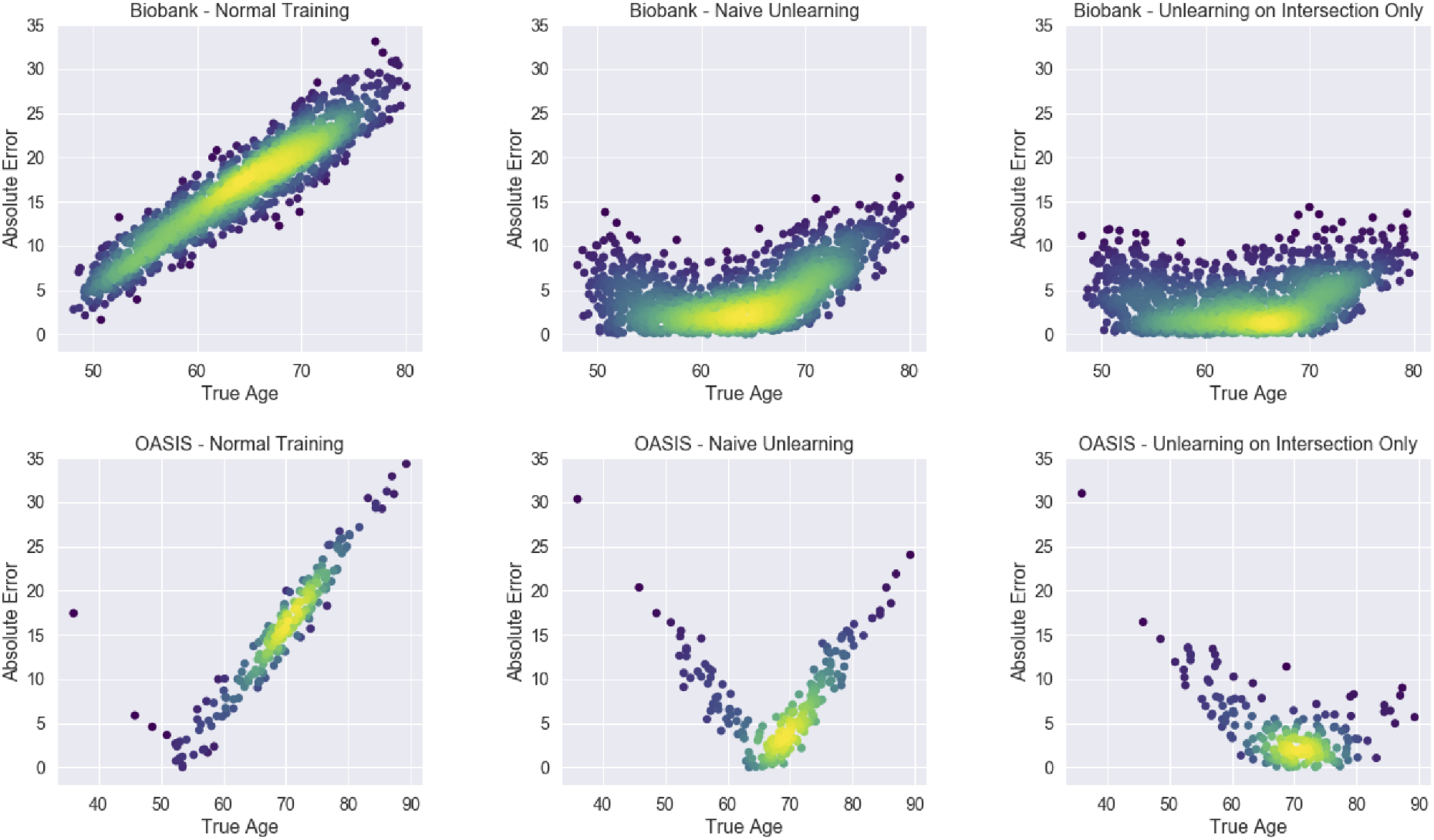
Density plots showing the absolute errors for the three different training regimes: standard training, naïve unlearning and unlearning only on the overlap data for 10-year overlap. It can be seen that unlearning only on the overlap dataset leads to substantially lower losses across both datasets.

As expected, it can be seen that the normal training regime produces large errors, especially outside of the range of the Biobank training data, as the learnt weights are very largely driven by the much larger size of the Biobank training data. With naïve unlearning, the network is not able to correct for both scanners and the results for the OASIS data are poor, whereas by unlearning on just the overlapping subjects, the error is reduced on both of the datasets during testing. The only area where we see a reduction in performance is the lower end of the OASIS dataset, possibly because when the network was being driven by the Biobank data, the network generalised well to the OASIS subjects from the same range. Naïve unlearning also performs slightly less well at removing scanner information on the testing data, probably indicating that the features removed also encode some age information and so generalise less well across the whole age range.

#### 4.1.3. Extension to ABIDE data

Figure 8 shows the results of applying the unlearning process to the ABIDE data, comparing normal training to unlearning. It can be seen that unlearning improves the MAE across all sites apart from MPG, which contains significantly older subjects compared to the other sites. The scanner classification accuracy before unlearning was 55.9% and after unlearning was reduced to 6.42%, where random chance was 6.25%. Therefore, we can see that scanner information is present in the feature embeddings, even when harmonised protocols are used.

**Figure 8:**
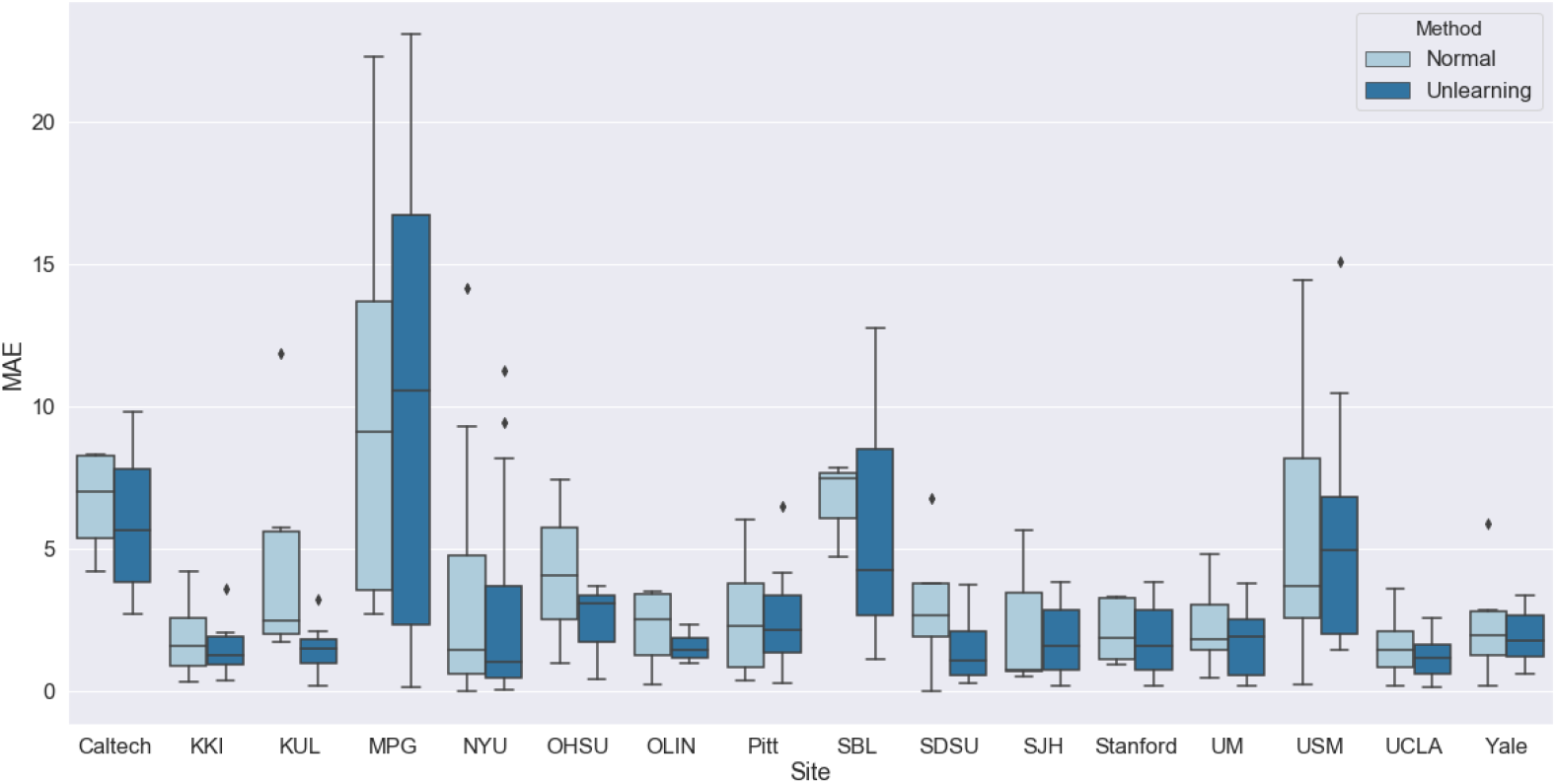
MAE results broken down by site for the ABIDE data, comparing normal training to unlearning.

These results therefore show that the framework can be applied to many sites with no changes needed. It also shows that the framework is applicable to lower numbers of subjects, with some of the sites having as few as 27 subjects for training. We thus foresee that the framework should be applicable to many studies.

#### 4.1.4. Removal of other categorical confounds

The effect of removing sex information as an additional confound in addition to harmonising for scanner was investigated. In Fig.9 it can be seen that there is no significant effect on the MAE results when removing sex information in addition to scanner information. Unlearning sex information had no substantive effect on the ability to remove scanner information with the scanner classification accuracy being 48% and 49%, respectively. The sex classification accuracy was 96% before unlearning and 54% after unlearning. Therefore, we can remove multiple pieces of information simultaneously and, so long as the information we wish to unlearn does not correlate with the main task, we can do so without significantly reducing the performance on the main task.

**Figure 9:**
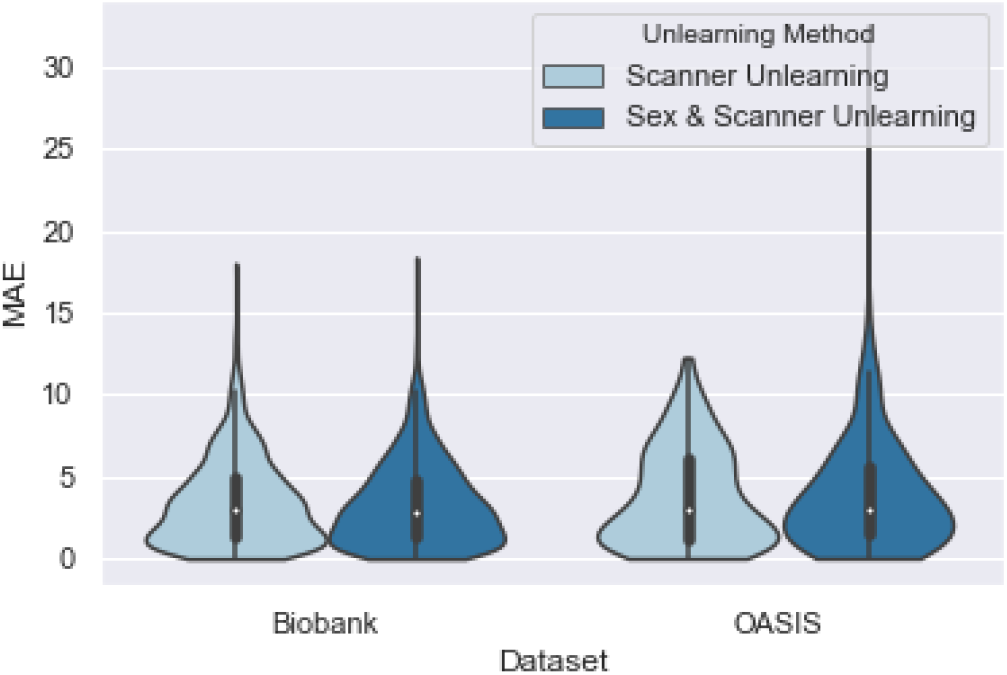
Results comparing unlearning scanner only to unlearning scanner and sex information.

We then considered the case where sex information was correlated with the acquisition scanner given that 80% of the subjects in the OASIS dataset are male and 80% of the subjects in the Biobank dataset are female. Table 4 shows the comparison of normal training on these datasets compared to unlearning sex on all data points and unlearning sex on a subset with balanced numbers of each sex for each scanner. The full testing set was still used.

**Table 4:**
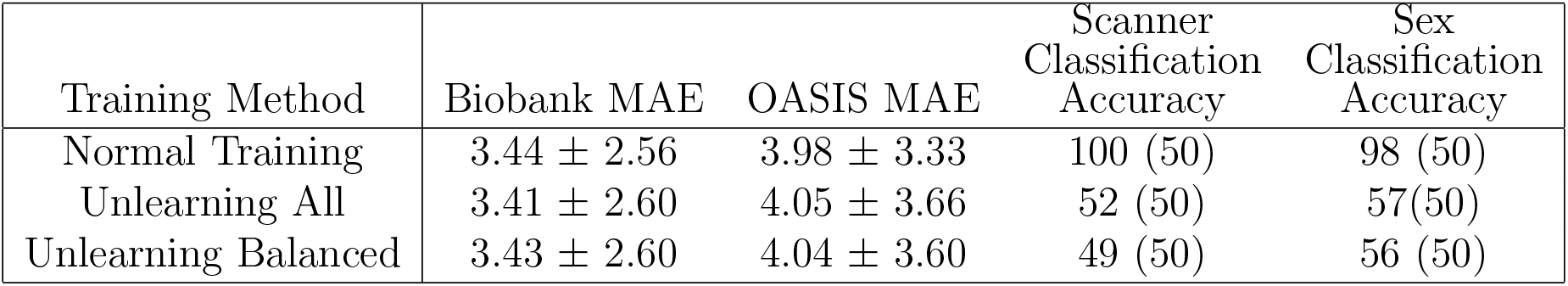
MAE results for unlearning scanner and sex when sex is highly correlated with scanner.

It can be seen that there is little difference between unlearning on all, or just a balanced subset, compared to the normal training baseline. It can also be seen that neither method affects the network’s ability to unlearn scanner or sex information and so there is no need to change the unlearning training from the standard case.

Finally, we considered the case where sex – the confound we wish to unlearn – is highly correlated with age – the main task – such that 80% of the subjects below the median age are female and 80% of the subjects over the median age are male. Again, normal training was compared to unlearning on the whole dataset and to unlearning on a curated subset, with uniform distributions of sex with age for each scanner. Testing was again performed on the full testing datasets.

It was found that it was not possible to unlearn on the whole distribution as it caused the training to become unstable almost immediately (after approximately 5 epochs), with the training and validation loss exploding before we were able to unlearn sex and scanner information. The results from unlearning scanner can be seen in Table 5 where it can be seen that by unlearning only on a subset of data points with equal sample numbers we can almost entirely unlearn both scanner and sex information. Figure 10 shows a T-SNE [31] of ***Q**_repr_* where it can be seen that, before unlearning, the data was largely separable by scanner and sex, and that, after unlearning, these are indistinguishable.

**Table 5:** MAE results for unlearning scanner and sex when sex is highly correlated with age.

| Training Method | Biobank MAE | OASIS MAE | Scanner<br>Classification<br>Accuracy | Sex<br>Classification<br>Accuracy % |
| --- | --- | --- | --- | --- |
| Normal Training | $3.45 \pm 2.60$ | $4.85 \pm 3.73$ | 100 (50) | 95 (50) |
| Unlearning All | $5.25 \pm 3.70$ | $5.62 \pm 4.07$ | 100 (50) | 65 (50) |
| Unlearning Balanced | $3.53 \pm 2.61$ | $4.53 \pm 3.52$ | 52 (50) | 54 (50) |

**Figure 10:**
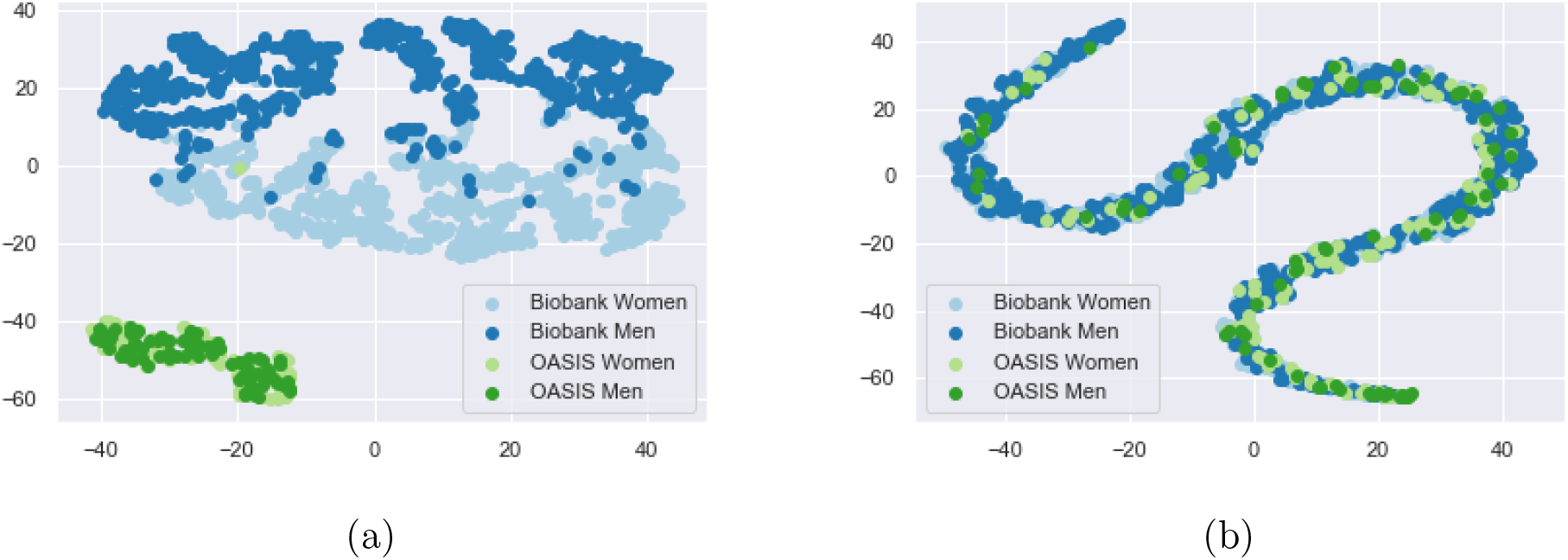
a) T-SNE plot of ***Q**_repr_* from training without unlearning, having trained on a dataset where sex correlates highly with age. It can be seen that the scanners can still be almost entirely separated, except for two data points grouped incorrectly (located within the cluster of light blue data points), showing that data from each scanner has its own distinct distribution and that the data points can also be entirely split by sex for the Biobank data. b) T-SNE plot of ***Q**_repr_* after unlearning. It can be seen that, through the unlearning, the distributions become entirely jointly embedded in terms of both scanner and sex.

#### 4.1.5. Removal of Continuous Confounds

For the removal of age as a confound, we used sex prediction as the main task. We achieved an average of 96.3% on the sex prediction task before unlearning and 95.9% after unlearning. As before, we were able to unlearn scanner information, reducing the scanner classification accuracy from 100% to 53%. Figure 11a) shows the averaged softmax labels from the age prediction task with normal training, where it can be seen that there is a large degree of agreement between the true labels and the predicted labels, showing that we are able to learn age using the continuous labels and KL divergence as the loss function. We achieved MAE values of 3.26 ± 2.47 for Biobank and 4.09 ± 3.46 for OASIS with normal training in this manner.

**Figure 11:**
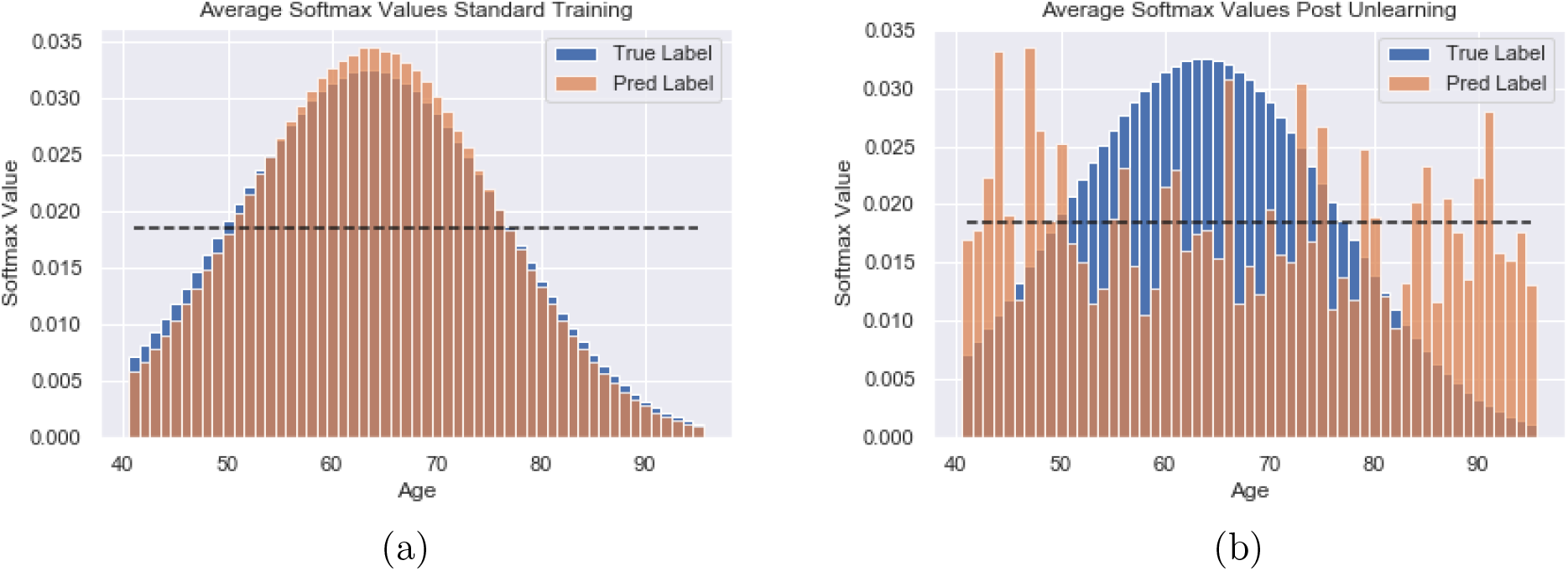
Softmax labels for the age prediction task, averaged across all examples for the Biobank dataset showing both the true values converted into Gaussian distributions and the predicted values. The dashed line corresponds to the value if all the softmax values were equal. a) Predictions with standard learning b) Predictions after unlearning. It can be seen that after unlearning the softmax values are much closer to the dashed line and there is no trend towards the true maximum.

Figure 11 b) shows the softmax labels after unlearning. It can be seen that the predicted labels no longer follow the same distribution as the true labels and that they are distributed around the random chance value indicated by the dotted line. It can also be seen that there is no trend towards the true age value, indicating that a large amount of the age information has been removed. After unlearning, the MAEs for the age prediction task were 17.54± 7.65 for Biobank and 22.86 ± 9.40 for OASIS. The average KL divergence has increased from 0.0022 with standard learning to 0.251 after unlearning. This therefore demonstrates that, by adapting the labels, we can also use the same training regime to remove continuous information.

### 4.2. Segmentation Task

#### 4.2.1. Supervised Learning

The results from comparing the location of the domain classifier can be seen in Table 6. We compare the Dice scores and consider the scanner classification accuracy. It can be seen that the best results for both Dice and the scanner classification accuracy were achieved with the domain classifier simply connected at the final convolution. Training with the domain classifier attached to the bottleneck was also much less stable. Given this result, the location of the domain classifier is fixed to the final convolutional layer for the rest of the experiments that follow.

**Table 6:** Dice scores comparing different locations for attaching the domain classifier in the network (as indicated in Fig.5) A) at the final convolutional layer, B) at the bottleneck, and A+B) the combination of the two locations. The scanner classification accuracy was the accuracy achieved by a separate domain classifier using the fixed feature representation at the final convolutional layer. Random chance is given in brackets.

| Location of<br>Domain Classifier | Biobank Dice | OASIS Dice | Scanner<br>Classification<br>Accuracy |
| --- | --- | --- | --- |
| Final Convolution (A) | $0.910 \pm 0.023$ | $0.916 \pm 0.021$ | 51 (50) |
| Bottleneck (B) | $0.871 \pm 0.046$ | $0.882 \pm 0.030$ | 100 (50) |
| Both (A+B) | $0.903 \pm 0.025$ | $0.912 \pm 0.021$ | 55 (50) |

This finding goes against the expectation from the literature [28] which suggests a mixture of locations are best for domain adaptation. We suspect this is due to our prioritisation of the removal of the scanner information to ensure harmonisation through unlearning: at the bottleneck (B) we have the most compressed representation of the data and it is probable that by unlearning at that location we constrain these features too highly and they are unable to find as successful a representation of the data.

The results for training on the combination of datasets with normal training and unlearning can be seen in Table 7, averaged across tissue type, and in Fig.12 by tissue type for each scanner. It can be seen that the unlearning process does not reduce the Dice score achieved across tissue types and that we are able to almost entirely unlearn scanner information.

**Table 7:**
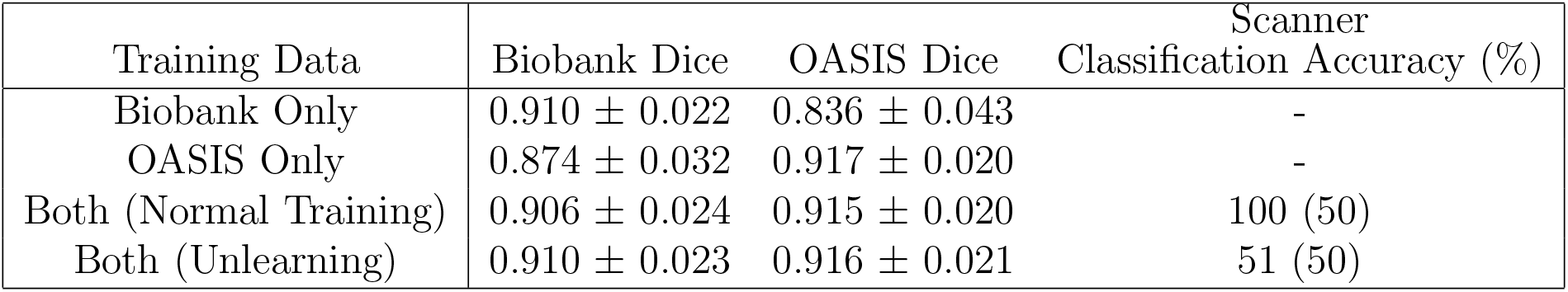
Dice scores comparing unlearning to training the network on different combinations of the datasets, averaged across the tissues types. Scanner accuracy is the accuracy achieved by a domain classifier given the fixed feature representation, with random chance given in brackets.

**Figure 12:**
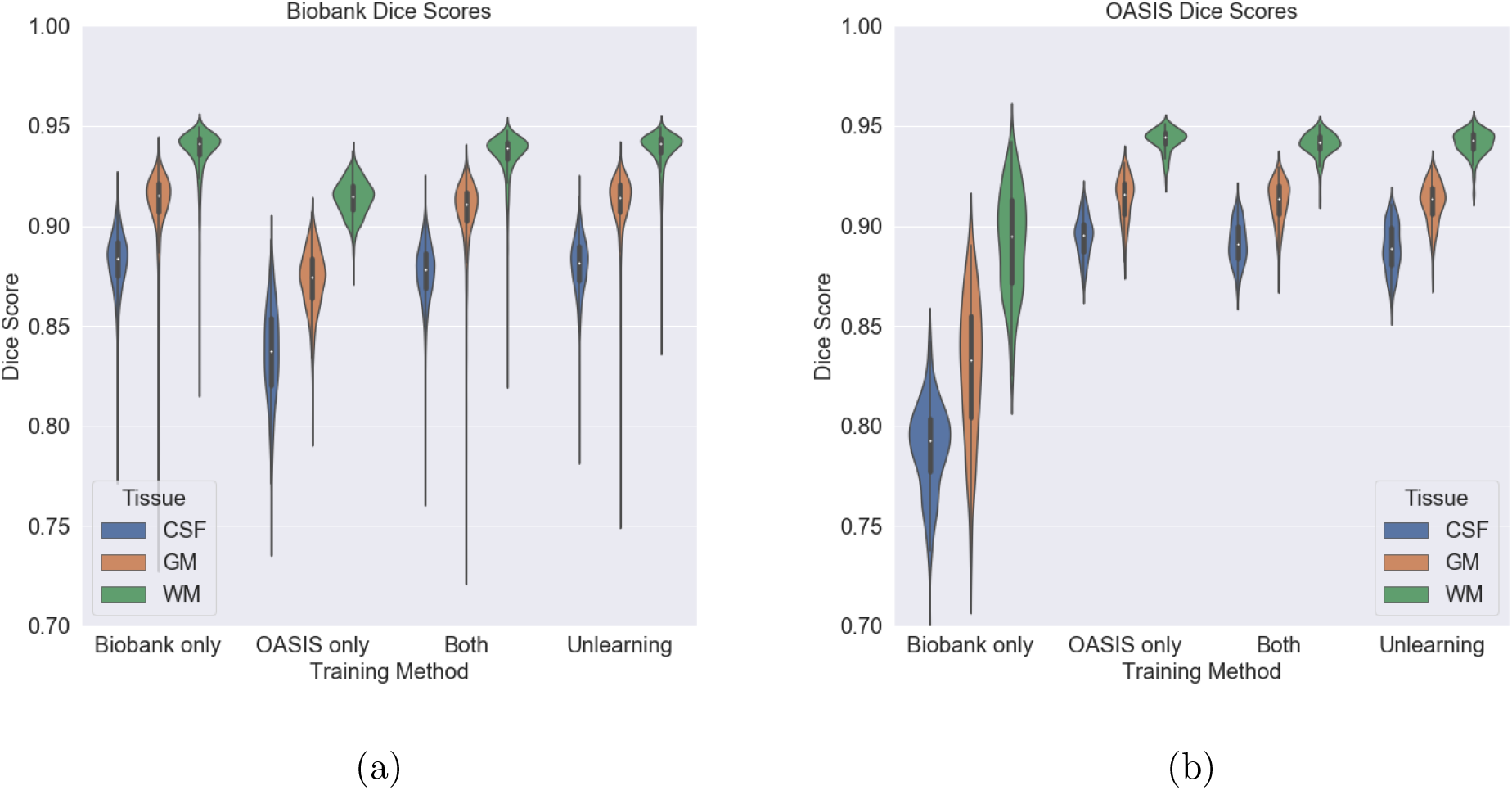
Dice scores for the two datasets for each method broken down by tissue type. CSF = Cerebrospinal fluid Fluid, WM = White Matter, GM = Grey Matter

#### 4.2.2. Semi-Supervised Segmentation

To allow us to explore the effect of training the network with low numbers of labelled training points for one of the scanners, we trained the network with both normal learning and with unlearning for different numbers of OASIS data points with labels. Only the examples with labels were used for evaluating the main task but all of the data points were still used for unlearning scanner as these loss functions do not require main task labels to be evaluated. The results can be seen in Fig.13: for all cases, the scanner classification accuracy for normal training was 100% and between 50% and 55% for unlearning. It can be seen that unlearning gives a large improvement in the segmentations with low numbers of data points, not only in terms of the mean value but also the consistency of the segmentations, including when the network is unsupervised with regards to the OASIS dataset and so has no training labels. Even with large numbers of training examples, there is never a disadvantage to using unlearning.

**Figure 13:**
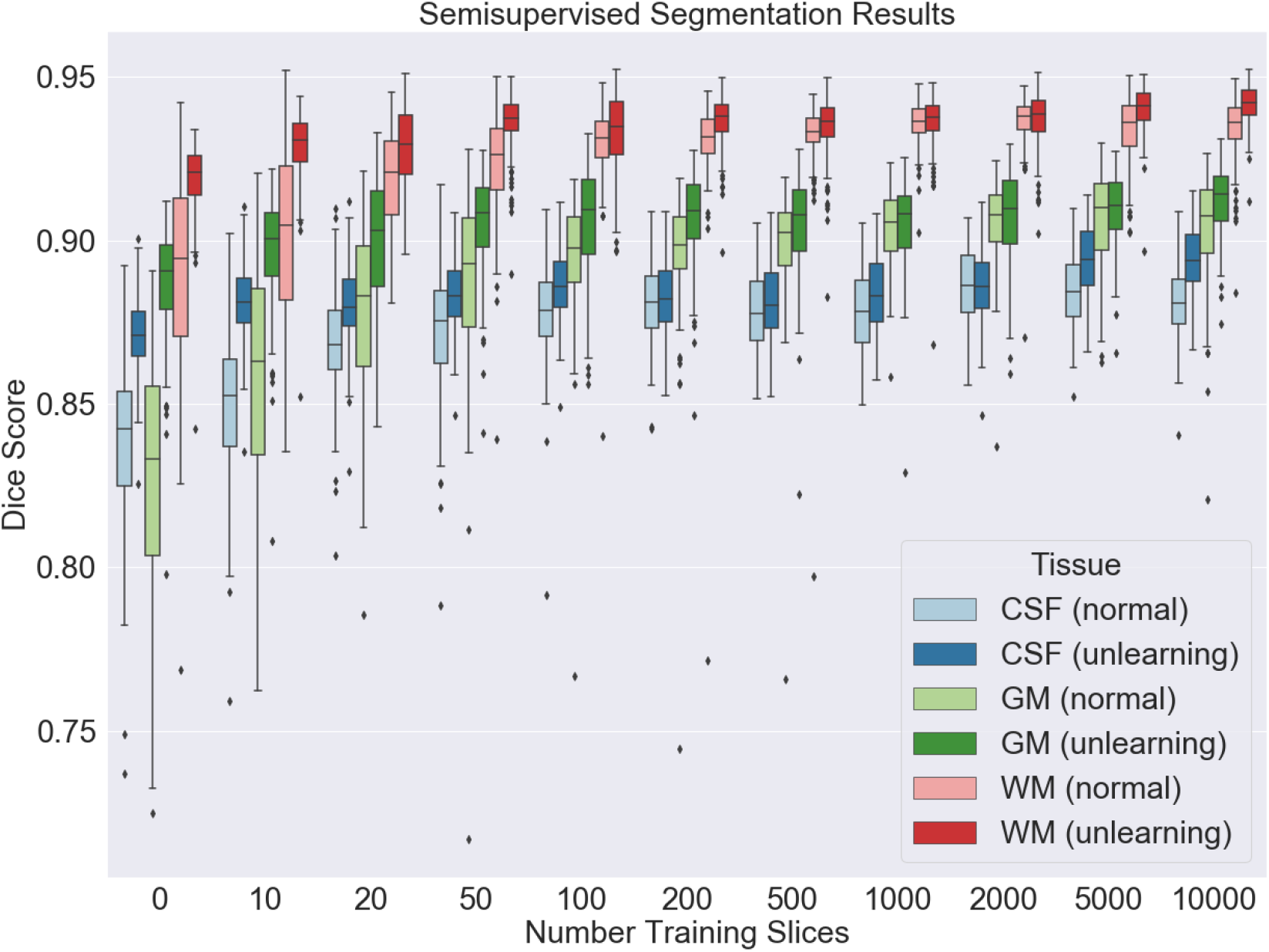
Dice scores for the three different tissue types for the OASIS data with increasing numbers of OASIS training slices, comparing both normal training and unlearning with the full Biobank dataset used throughout. For clarity, the x axis is not plotted to scale.

**Figure 14:**
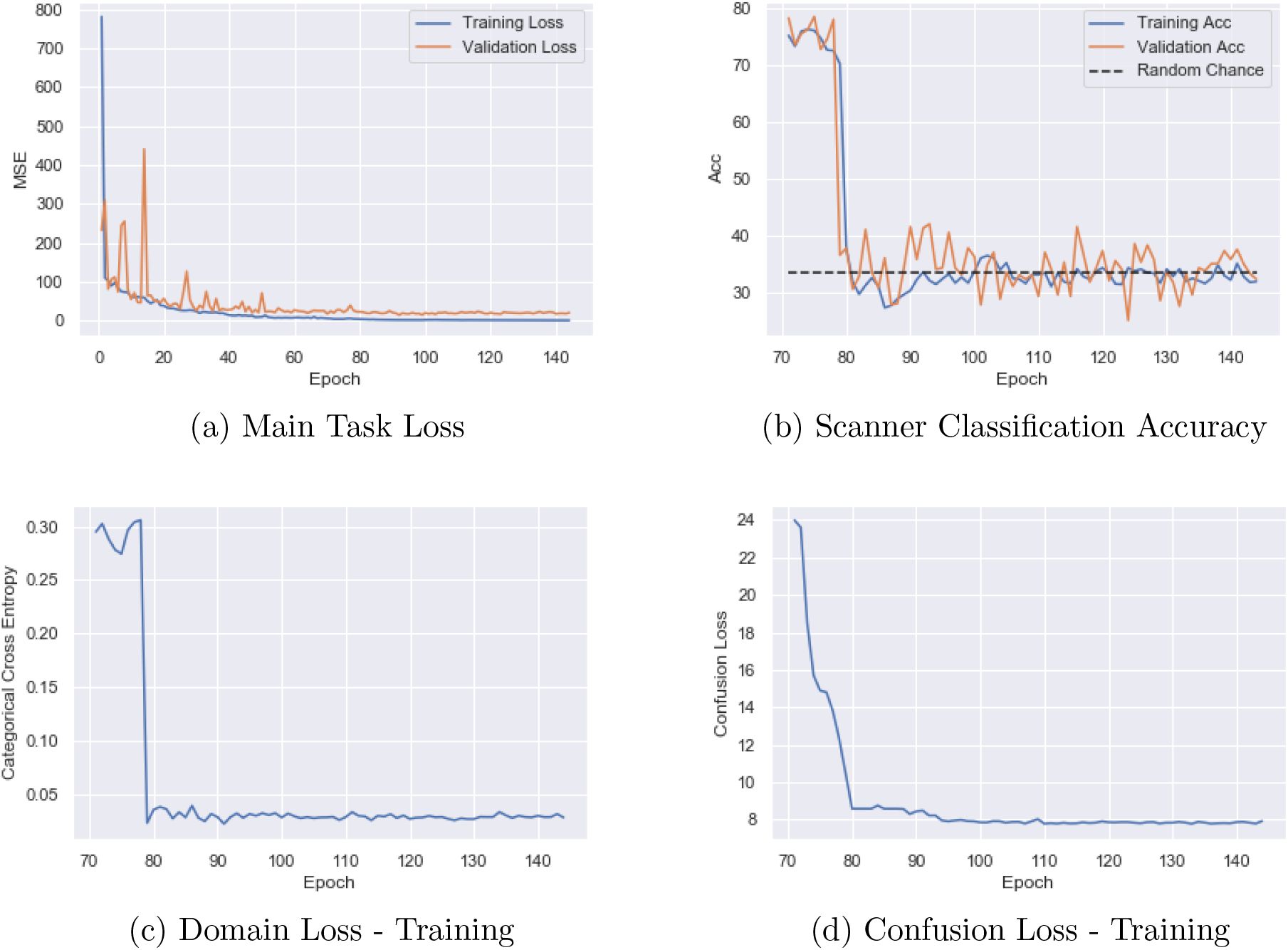
Graphs for the different loss functions from training. Only the main task loss is used during pretraining and so all other losses begin at 70 epochs when convergence on the main task is achieved.

## 5. Discussion

In this work, we have shown how a domain adaptation technique can be adapted to harmonise MRI data for a given task using CNNs, by creating a feature space that is invariant to the acquisition scanner. Removal of scanner information from the feature space prevents the information driving the prediction. The cost of the harmonisation is an approximately two fold increase in training time (from 0.88 seconds per batch to 1.53 seconds) but there is no increase in prediction time, and so, once the model is trained, there is no disadvantage to having unlearned scanner information. The duration of a batch increase is not the three times that might be expected, with the increase from one-forward and backward pass to three forward and backward passes per epoch, because the second two loss functions only update subsets of the parameters, reducing the number of calculations that must be completed.

We have demonstrated our technique for both regression and segmentation and so it should be applicable to nearly all medical imaging problems. The applicability to a wide range of networks, even those with skip connections such as the U-Net, provides a high level of flexibility. It does, however, only harmonise the features for a given task and therefore a separate network must be trained for each task. Consequently, an area for future research would be to harmonise across both scanners and tasks. As a comparison, a DANN-style network [18], with the same architecture as our network but an additional gradient reversal layer inserted for the domain classifier branch, was also trained, but it was found that when training with only two datasets, the domain classifier was unstable and predicted the domains with 7.2% accuracy (results can be seen in the supplementary material). This was a local minimum but clearly meant that the scanner information was not successfully removed. This instability in adversarial tasks posed as a minmax optimisation has also been experienced for other applications across the literature [30, 21]. By contrast, the confusion loss used in our iterative approach forces the logits towards random chance and we experience much more stability in the domain unlearning branch. We have not been able to compare to existing harmonisation methods, such as ComBat, because the different approaches are not easily compared, especially for the segmentation task, and focus on different stages of the pipeline.

A potential limitation of our work is that the framework is not generative and therefore cannot be easily used in conjunction with tools such as VBM or Freesurfer, although could be added into deep learning equivalents such as Fastsurfer [23], especially as more tools gain CNN-based equivalents. The decision to not make the framework generative was due to generative CNN methods needing large amounts of data to train or, even in some examples, paired data, meaning that they are not applicable to most real life neuroimage studies, whereas we have shown that our framework is applicable even on highly multi-site datasets with limited examples from each site, such as the ABIDE data. Furthermore, due to the data-hungry nature of generative CNNs, most of these methods reduce 3D volumes to 2D slices or even patches. This then leads to errors when reconstructing the outputs, such as inconsistencies between slices. Were these images then to be used in further downstream analysis, it is very hard to anticipate how these errors might impact on obtained results and to know how to account for them. Although we do not produce harmonized images, the image-derived values can be used in downstream analysis, for instance tissue volumes or segmentation maps can be used with a GLM to explore group differences. Our approach, therefore, is limited to within CNNs but this is a rapidly growing area of study within neuroimaging and the flexible nature of the framework means it should be applicable across feedfoward architectures and different neuroimaging tasks.

The ablation study on the age prediction task shows that not only are we able to unlearn the scanner information but that there is an increase in performance that is not just due to the main task loss function. This indicates that the learned features generalise better across datasets, which is confirmed by the performance on the third dataset when only training on two. This corroborates the scanner classification accuracy and T-SNE plots, showing that scanner information has largely been removed and that the features that remain encode age information that transfers across datasets. Even for the segmentation task, where there is no significant improvement in the performance on the segmentation task, including the unlearning gives us the assurance that the output values are not being driven by scanner. This could be important for downstream analysis, for instance comparing white matter volumes, allowing the data to be pooled across sites. This result is also robust to the choice of the weighting factors and so the unlearning is not sensitive to the hyper-parameters.

The results also show that we can use the unlearning scheme even when there is a strong relationship between the main task label and the acquisition scanner. This could be especially useful when combining data between studies with different designs and with very different numbers of subjects from each group. As we can perform unlearning on a different dataset to the main task, we have the flexibility to apply the method to a range of scenarios.

At the limit where there exists no overlap between the datasets’ distributions, unlearning scanner information would be highly likely to remove information relating to the main task. This would then only be able to be solved by having an additional dataset for each scanner, acquired with the same protocol, and so represents a potential limitation of the method. The extension to the ABIDE dataset, however, shows the applicability of the method to many small datasets, and therefore this may be solvable using a set of small datasets which together span the whole distribution. It also shows that the framework is able to harmonise across scanner manufacturers as the data was collected on a range of GE, Siemens and Philips scanners.

We have also shown that the approach can be extended to allow us to remove additional confounds, and have demonstrated a way that this could also be extended to allow us to remove continuous confounds such as age. For each confound to be removed, we require two additional loss functions and two additional forward and backward passes. Therefore, the training time will increase with each additional confound, presenting a potential limitation. Were many confounds to be removed, we might also need to increase the number of times the passes for the main task are performed to prevent the performance on the main task suffering too much degradation due to the feature space being optimised for multiple tasks. We have, however, also shown how confounds can be removed even when they correlate with the main task. By carefully selecting a subset of the data with which to unlearn the scanner, we can remove the confounds, including, for instance, where sex is correlated with age, which is a case where both the main task and confound to be removed are associated with structural changes.

The approach can also be applied when no labels are available for one or more of the domains. We have demonstrated for the segmentation task that we are able to use this technique effectively when we only have labels for one domain and we would expect that this should extend to multiple unlabelled datasets.

## 6. Conclusion

We have presented a method for MRI harmonisation and confound removal that should be applicable across many tasks for neuroimaging and data scenarios. We have shown that it can be easily applied to segmentation, classification and regression tasks and with the highly flexible nature of the framework it should be applicable to any feedforward network. The ability to remove scanner bias influencing the predictions of CNNs should enable to the combination of small datasets and the exploration of problems for which there are no single-scanner datasets of adequate size to consider.

## 7. Data and Code Availability Statement

The data used in these experiments are available on application to the relevant studies. The code used is available at www.github.com/nkdinsdale/Unlearning_for_MRI_harmonisation and weights from training are available on request through emailing the corresponding author.

## 8. Acknowledgements

ND is supported by the Engineering and Physical Sciences Research Council (EPSRC) and Medical Research Council (MRC) [grant number EP/L016052/1]. MJ is supported by the National Institute for Health Research (NIHR), Oxford Biomedical Research Centre (BRC), and this research was funded by the Wellcome Trust [215573/Z/19/Z]. The Wellcome Centre for Integrative Neuroimaging is supported by core funding from the Wellcome Trust [203139/Z/16/Z]. AN is grateful for support from the UK Royal Academy of Engineering under the Engineering for Development Research Fellowships scheme.

This research has been conducted in part using the UK Biobank Resource under Application Number 8107. We are grateful to UK Biobank for making the data available, and to all UK Biobank study participants, who generously donated their time to make this resource possible. Analysis was carried out on the clusters at the Oxford Biomedical Research Computing (BMRC) facility and FMRIB (part of the Wellcome Centre for Integrative Neuroimaging). BMRC is a joint development between the Wellcome Centre for Human Genetics and the Big Data Institute, supported by Health Data Research UK and the NIHR Oxford Biomedical Research Centre.

The computational aspects of this research were supported by the Wellcome Trust Core Award [Grant Number 203141/Z/16/Z] and the NIHR Oxford BRC. The views expressed are those of the author(s) and not necessarily those of the NHS, the NIHR or the Department of Health.

The primary support for the ABIDE dataset by Adriana Di Martino was provided by the (NIMH K23MH087770) and the Leon Levy Foundation. Primary support for the work by Michael P. Milham and the INDI team was provided by gifts from Joseph P. Healy and the Stavros Niarchos Foundation to the Child Mind Institute, as well as by an NIMH award to MPM ( NIMH R03MH096321).

## 10. Supplementary Materials

### 10.1. Training Graphs

Example training graphs from the age prediction task with three datasets. Only the main task loss (MSE for age prediction) is used during pretraining and so the other loss functions begin at epoch 70 when unlearning started. Unlearning began when convergence was achieved on the main task (patience = 15 epochs). It can be seen that beginning unlearning does not reduce the performance on the main task. It can also be seen that all the loss functions are stable throughout training.

### 10.2. DANN Results

A DANN-style domain adaptation approach was also tried due to its simpler implementation. As confirmed across the literature, it was unstable to train, limiting its performance. The results from training can be see in Table 8 where it can be seen that the scanner classification accuracy was much lower than random chance, indicating that scanner information remained in the feature representation. It can also be seen that the performance on the main task was reduced, with the domain adaptation leading to lower performance compared to normal training.

**Table 8:**
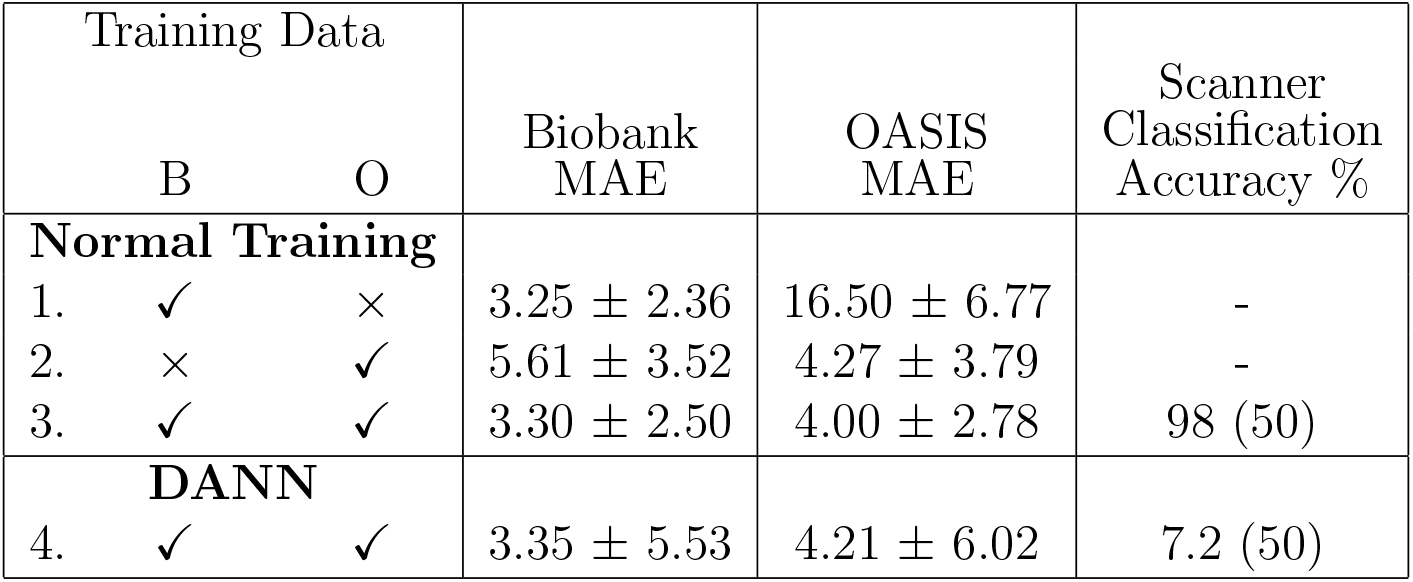
Comparing normal training on the different combinations of datasets to using DANN to remove scanner information.

### 10.3. Effect of Hyper-Parameters

For all experiments, the value of *α* remained set to 1 and no change was required. The value of *β* was varied between experiments, controlling the training stability and the rate of convergence: higher β values provide more stability for the domain unlearning but decrease the rate of convergence for the main task. It should not have any effect on the final age prediction values achieved, given that for all values we are able to reach convergence, and so we varied the value of β between 0.1 and 1000 and compared the MAEs for the three datasets. It can be seen that the value of *β* has no effect on the achieved MAEs across these values from Fig.15. Therefore, the segmentation performance is robust to the choice of β and the value chosen can be selected to maximise the stability of training without impacting on the final prediction values. Stability could also be controlled by using different learning rates for each stage; however, this was found to be harder to tune in practice.

**Figure 15:**
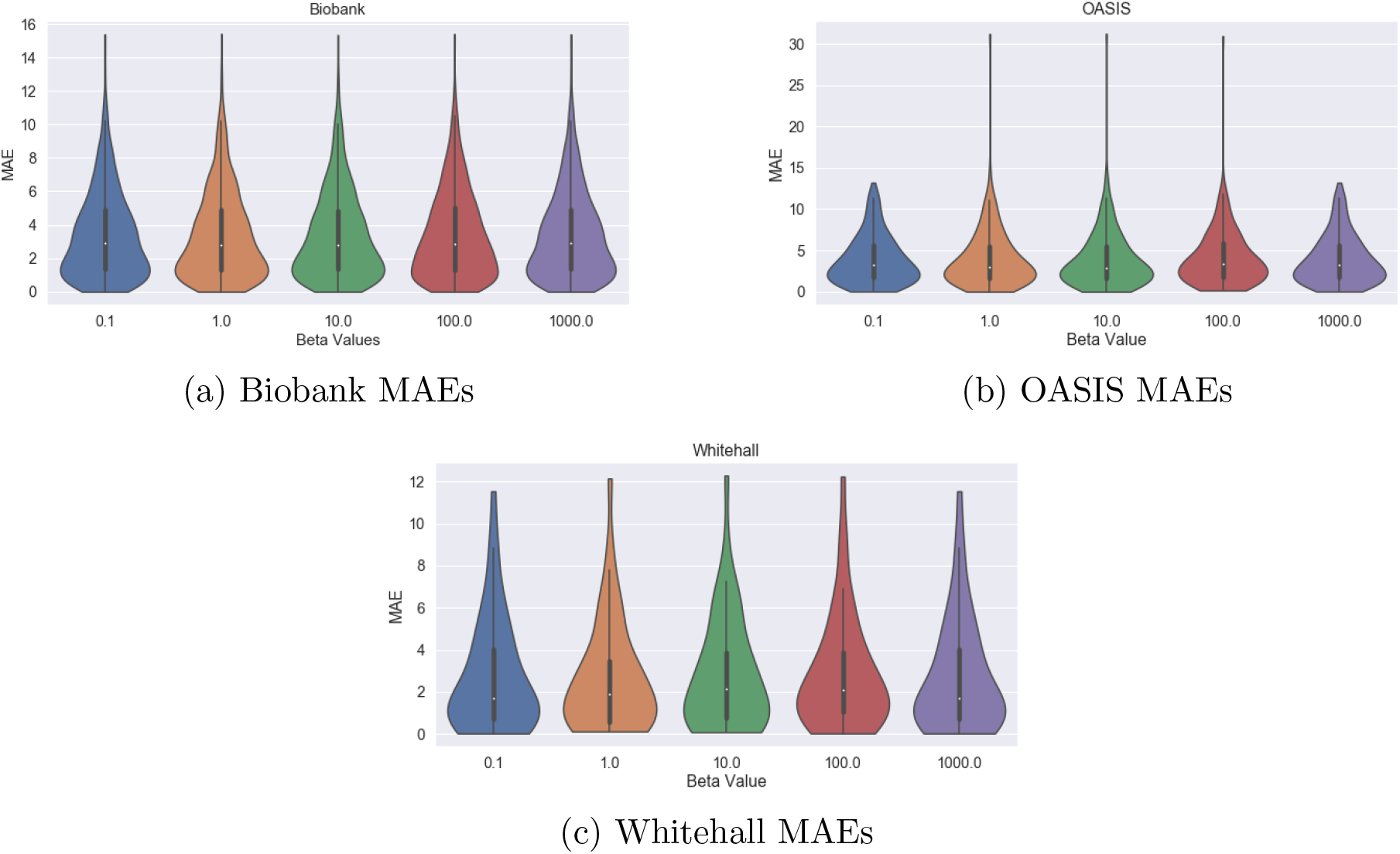
MAEs for each dataset with increasing values of the constant *β* used in training, where *β* is the weight determining the contribution of the confusion loss.

Finally, we explored the choice of the batch size used. For training, the largest batch that could fit into memory was used to obtain the results in the paper; here the effect of smaller batches is explored. Figure 16 shows the validation loss during training for batch sizes of 3, 8 and 32 (other batch sizes are not shown for clarity) and the pattern was the same across the different loss functions. It can be seen that the training was more stable with the smaller batch sizes. However, it can be seen from Fig.17 that the overall MAEs achieved across the datasets were better with larger batch sizes. This is most likely because larger batch sizes force the network to unlearn scanner information more thoroughly than, for instance, with a batch size of 3, where given our constraint on the batches, there can only be one example per scanner. Given the result from varying the value of *β* above, we should select the batch size so as to maximise the performance on the main task and vary the β value to control the stability of training.

**Figure 16:**
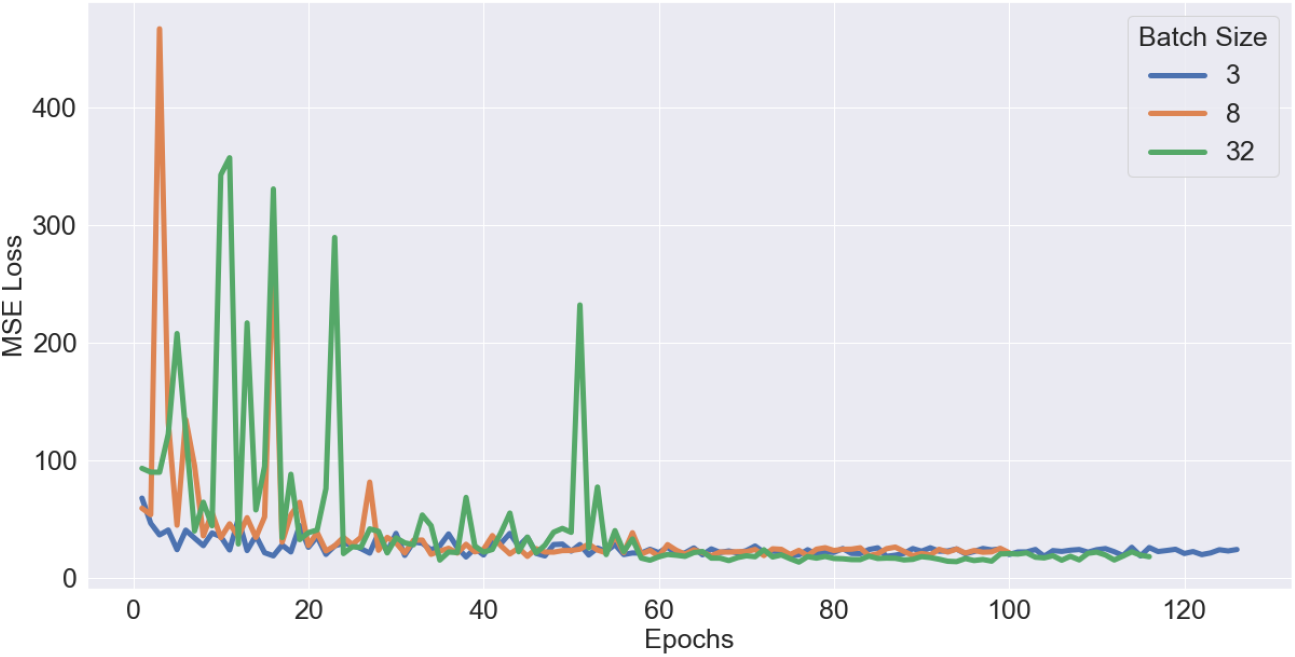
Validation loss with epochs for batch sizes of 3, 8 and 32.

**Figure 17:**
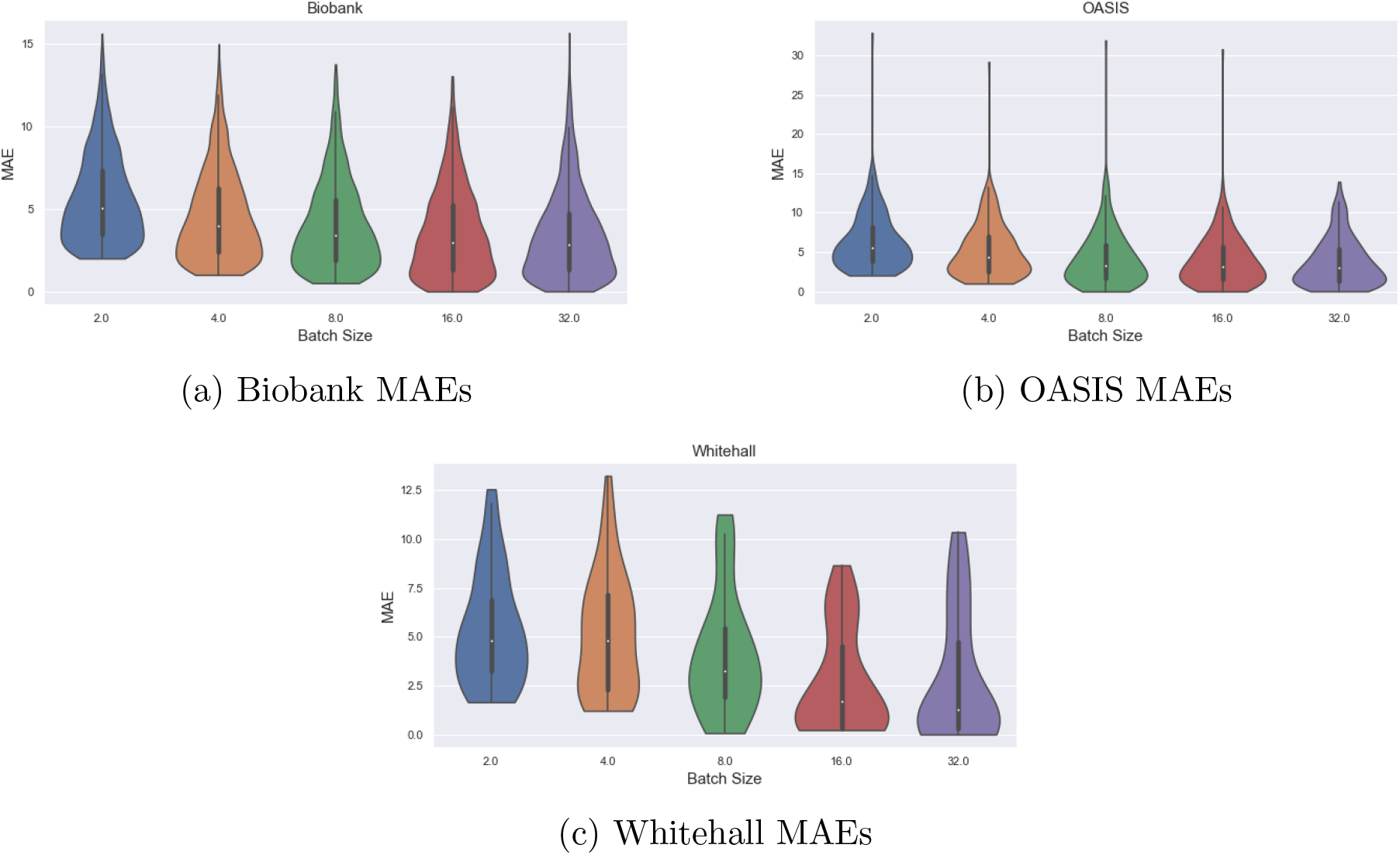
MAEs for each dataset with increasing batch sizes used during training.

**Figure 18:**
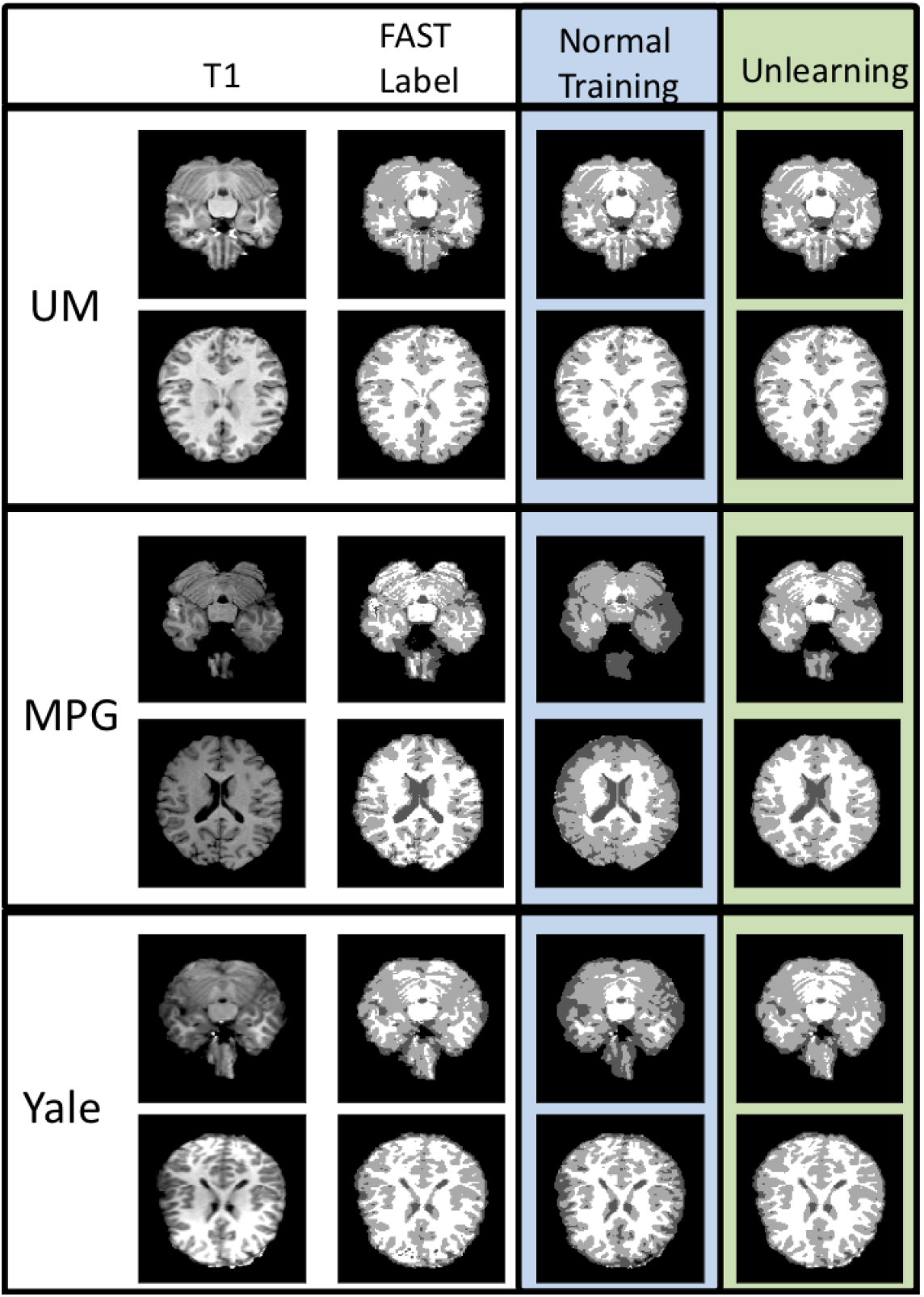
Segmentation results for the ABIDE data using normal training and unlearning when labels are only available for the UM site. Labels are white matter, grey matter and CSF.

**Figure 19:**
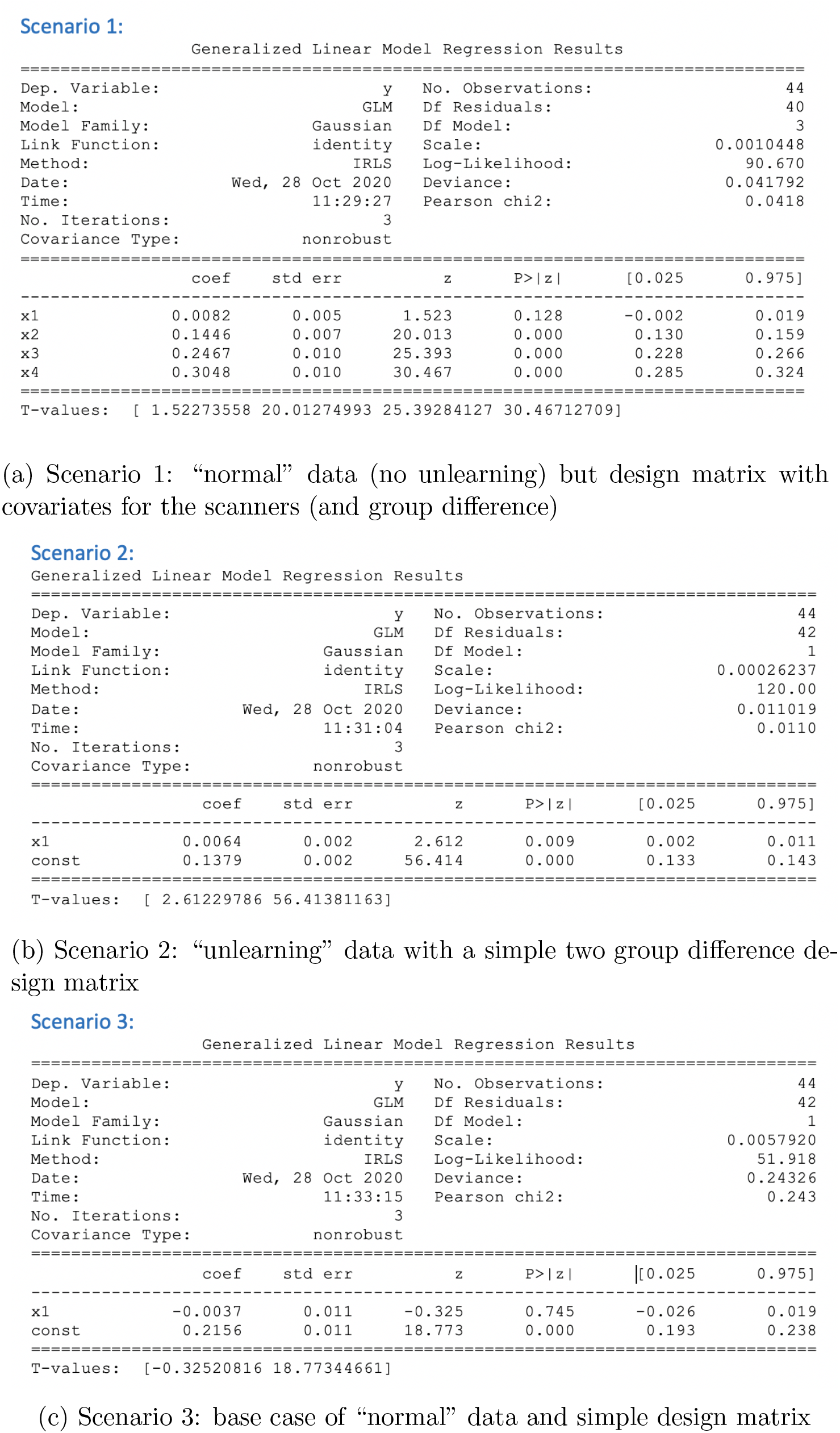
GLM results for the three explored scenarios.

### 10.4. p Values

Paired t-tests have been completed between experiments and the uncorrected p-values are reported separately for each dataset. For each table in the results section, the rows compared are indicated in the first column of the table.

#### 10.4.1. Age Prediction - Basic Fully-Supervised Learning

| Rows Compared | Biobank | OASIS | Whitehall |
| --- | --- | --- | --- |
| 4) - 9) | 0.038 | 0.567 | 0.621 |
| 5) - 10) | 0.094 | $1.63 \times 10^{-14}$ | 0.001 |
| 6) - 11) | $1.69 \times 10^{-14}$ | 0.001 | $1.21 \times 10^{-10}$ |
| 7) - 12) | 0.002 | 0.042 | $4.61 \times 10^{-7}$ |
| 4) - 13) | 0.223 | 0.923 | 0.032 |
| 5) - 14) | 0.099 | $9.42 \times 10^{-15}$ | $8.62 \times 10^{-20}$ |
| 6) - 15) | $4.21 \times 10^{-23}$ | 0.012 | 0.008 |
| 7) - 16) | 0.145 | $3.25 \times 10^{-12}$ | $1.96 \times 10^{-22}$ |
| 9) - 13) | 0.416 | 0.004 | 0.070 |
| 10) - 14) | 0.893 | 0.034 | 0.041 |
| 11) - 15) | $3.96 \times 10^{-24}$ | $1.30 \times 10^{-4}$ | $3.61 \times 10^{-8}$ |
| 12) - 16) | 0.011 | 0.666 | 0.098 |

- 1) - 16) Biobank Only: 0.103 2) - 16) OASIS Only: $5.62 \times 10^{-31}$ 3) - 16) Whitehall Only: $9.71 \times 10^{-71}$

#### 10.4.2. Age Prediction - Balanced Datasets

| Rows Compared | Biobank | OASIS | Whitehall |
| --- | --- | --- | --- |
| 1) - 3) | $4.65 \times 10^{-11}$ | $2.34 \times 10^{-18}$ | $2.98 \times 10^{-10}$ |
| 2) - 4) | $5.67 \times 10^{-4}$ | $1.98 \times 10^{-3}$ | $7.45 \times 10^{-6}$ |

#### 10.4.3. Age Prediction - Biased Datasets10.4.4. Age Prediction - Sex Correlated with Scanner

10.4.3. *Age Prediction - Biased Datasets*
| Rows Compared | Biobank | OASIS |
| --- | --- | --- |
| 1) - 2) | $5.68 \times 10^{-89}$ | $8.62 \times 10^{-89}$ |
| 1) - 3) | $2.98 \times 10^{-92}$ | $5.78 \times 10^{-93}$ |
| 2) - 3) | $5.63 \times 10^{-4}$ | 0.098 |
| 4) - 5) | $3.69 \times 10^{-65}$ | $7.14 \times 10^{-75}$ |
| 4) - 6) | $3.48 \times 10^{-72}$ | $9.01 \times 10^{-83}$ |
| 5) - 6) | $9.72 \times 10^{-4}$ | 0.032 |
| 7) - 8) | $6.91 \times 10^{-42}$ | $4.78 \times 10^{-61}$ |
| 7) - 9) | $9.01 \times 10^{-51}$ | $1.00 \times 10^{-63}$ |
| 8) - 9) | 0.045 | 0.029 |

*10.4.4. Age Prediction - Sex Correlated with Scanner*

#### 10.4.4. Age Prediction - Sex Correlated with Scanner

| Rows Compared | Biobank | OASIS |
| --- | --- | --- |
| 1) - 2) | 0.456 | 0.892 |
| 1) - 3) | 0.416 | 0.493 |

*10.4.5. Age Prediction - Sex Correlated with Age*

#### 10.4.5. Age Prediction - Sex Correlated with Age

| Rows Compared | Biobank | OASIS |
| --- | --- | --- |
| 1) - 2) | 0.001 | 0.009 |
| 1) - 3) | 0.752 | 0.042 |

*10.4.6. Segmentation - Location of Domain Classifier*

#### 10.4.6. Segmentation - Location of Domain Classifier

| Rows Compared | Biobank | OASIS |
| --- | --- | --- |
| 1) - 2) | $9.72 \times 10^{-4}$ | $3.00 \times 10^{-3}$ |
| 1) - 3) | 0.003 | 0.091 |

*10.4.7. Segmentation - Method Comparison*

#### 10.4.7. Segmentation - Method Comparison

| Rows Compared | Biobank | OASIS |
| --- | --- | --- |
| 3) - 4) | 0.645 | 0.986 |

1) - 4) Biobank Only: 0.833 2) - 4) OASIS Only: 0.654

### 10.5. Comparison to Adding Site as Regressor

Here we explored whether there were an advantage to harmonising the network compared to training a network normally and adding scanner covariates to a linear model that is used for the analysis.

To do this we considered the situation where we have data from three sites and scanners but only have manual segmentation results for a single site, which is a common scenario when working with neuroimaging data as labels are expensive to acquire. We then wanted to train a network to complete the segmentation for our test subjects from all three sites (e.g. from the ABIDE data) and then use them in a standard GLM to explore the task of interest. In this case we considered exploring the relationship between age and the normalised grey matter/white matter/CSF volumes.

We trained the model using normal training on just the labelled data (UM) and also with unsupervised unlearning, harmonising the data (main task labels for just UM but unlearning scanner information from all three sites). When we trained the model on just UM and created output predictions for the remaining two sites, we found that we had good predictions for the test data from UM but there was a significant performance drop when we applied the method to the other two sites, due to domain shift. It can be seen from the figure below that although the network trained with normal training performs well on UM (average dice = 0.891 ± 0.015) the model suffers significant performance degradation when applied to the unseen sites – MPG (average dice = 0.689 ± 0.020) and Yale (average dice = 0.627 ± 0.039). It can also be seen that this loss in performance leads to significant failures such that image-derived values such as white matter or grey matter volumes are going to be very different from the expected values. Conversely, it can be seen that the performance is greatly improved using the harmonisation method even though no manual segmentation labels for MPG or Yale were used during training: average dice = 0.891 ± 0.019 (p=4e-8) and 0.892 ± 0.009 (p=4.5e-10), respectively. This comes at little cost to the performance of the segmentation on UM (average dice = 0.883 ± 0.03, p=0.015). Clearly a covariate approach would be unable to substantially reduce these image-level errors.

To go further and verify the effect for statistical tests on derived scalar quantities, we calculated the CSF volume, normalised by total brain volume. As we expect this to be indicative of age, we split the subjects into a young group and an old group, (n=44) and we used this in a GLM to explore the ability to find group differences. We considered three scenarios:

1. “normal” input data (no unlearning) but with a design matrix that has separate covariates for the scanners (as well as a simple two group, unpaired, difference regressor - this is x1 below),
2. “unlearning” data with a simple two group (unpaired) difference design matrix,
3. base case of “normal” data and simple design matrix with no scanner-based covariates. We compute this using the statsmodel python package. The results can be found in the figures below.

It can be seen that, as expected from the image examples, when we do nothing to correct for the scanner differences (scenario 3) we do not get a significant result (p=0.745). Equally, when we add scanner as a regressor in the model (scenario 1) we also do not get a significant result (p=0.128) but when we use our deep learning model we get a much stronger statistic and a significant difference (p=0.009).

Therefore, there are clear scenarios in which just adding scanner-based covariates in the model leads to reduced statistical differences, in this case preventing us achieving significant results. Consequently, there is a real need for harmonisation approaches, such as this unlearning method, in situations like this.

## Notes

https://github.com/nkdinsdale/Unlearning_for_MRI_harmonisation

### Competing Interest Statement

The authors have declared no competing interest.

